# Species-rich and genomically diverse: comparative genomics reveals how fusions, fissions, and sex chromosomes have shaped beetle evolution

**DOI:** 10.64898/2026.05.15.725483

**Authors:** Dwayne Tally, Charlie Pittman, Ryan R Bracewell

## Abstract

Chromosome evolution in animals reflects a balance between long-term conservation of ancestral linkage groups and lineage-specific chromosomal rearrangements that reshape genome structure. Beetles (Coleoptera), the most species-rich animal order, exhibit extensive diversity in karyotype, yet the extent to which their chromosomes retain deep ancestral structure remains unclear. Here, we analyzed 190 chromosome-level genome assemblies spanning 39 families and 16 superfamilies to characterize genome diversity, evaluate the conservation of ancestral linkage groups (Stevens elements), identify neo-sex chromosomes, and explore the role of repetitive elements in driving karyotypic change. Our results reveal that beetle genomes are highly diverse, varying substantially in genome size, chromosome number, GC content, and transposable element (TE) composition. Despite this diversity, Stevens elements appear conserved across much of the radiation, with several superfamilies maintaining strong chromosome synteny over more than 200 million years of evolution. In contrast, some clades, and specifically the leaf beetles (Chrysomelidae), have undergone extensive genomic changes including numerous chromosomal fusions and fissions and changes in genome size. Using synteny based approaches across beetles, we identified 37 species (approximately 19.5%) having patterns consistent with neo-sex chromosomes, a substantially higher frequency than previous estimates. These putative X-autosome fusions vary in complexity and age, often clustered within lineages that are prone to chromosomal instability. The ancestral X chromosome appears conserved for over 300 million years, with stable gene content and reduced TEs relative to autosomes. These findings help establish beetles as a promising system to uncover the evolutionary forces that maintain and disrupt ancestral linkage groups and drive the formation of neo-sex chromosomes.

## INTRODUCTION

Chromosomes are fundamental to the transmission of genetic material across generations. Many animal lineages exhibit deep conservation of chromosome-scale gene linkage, with entire chromosomes or ancestral linkage groups remaining intact over tens to hundreds of millions of years [1, 2]. However, rates of chromosomal evolution vary widely, with some groups undergoing frequent fusions, fissions, and inversions, while others maintain stable karyotypes despite deep evolutionary divergence [3]. Explaining why lineages differ so markedly in chromosomal stability is central to understanding constraints on genome architecture and the role of structural variation in diversification. Such variation likely reflects differences in forces acting on chromosomes, including variable mutation rates, recombination landscapes, repeat content, and effective population size, highlighting the importance of comparative genomic approaches across diverse taxa for disentangling the causes and consequences of genome restructuring over evolutionary time [4].

Among invertebrates, there is now substantial evidence that chromosomal architecture can remain stable over vast evolutionary timescales, in some cases persisting for hundreds of millions of years. In nematodes, this stability is exemplified by so-called “Nigon elements,” which represent conserved ancestral linkage groups that have been retained in many species despite deep evolutionary divergence [5]. Similarly, in insects, long-term chromosomal conservation has been documented in several holometabolous orders. For example, in flies (Diptera), “Muller elements” define deeply conserved chromosomal segments [6], while in butterflies and moths (Lepidoptera), “Merian elements” serve as conserved linkage groups [7]. Together, these findings highlight that despite ongoing evolutionary processes, large-scale genome organization in some invertebrate lineages can remain conserved over deep evolutionary time.

Beetles (Coleoptera) are the most species-rich group of animals, comprising more than 400,000 described species and representing approximately one quarter of all known animal diversity [8]. Despite longstanding interest in this extraordinary insect radiation (e.g., [9–13]), beetle genomes remain comparatively understudied relative to their diversity [14–16], and no comprehensive analysis of chromosome-scale genome structure has been conducted across the order, to date. Recent work identified a set of broadly conserved chromosomal blocks, termed Stevens elements, that correspond to ancestral linkage groups shared across beetle lineages [14]. Stevens elements were defined using the chromosomes of the model beetle *Tribolium castaneum*, analogous to the identification of Muller elements in Diptera [6, 17], and suggested substantial chromosomal conservation despite deep divergence among species. However, because analyses were limited by sparse taxonomic sampling and few chromosome-level assemblies, the extent to which Stevens elements are conserved across the beetle radiation remains unclear. As a result, broad patterns of genome evolution and structural diversity across Coleoptera are still poorly resolved, and heterogeneous annotation pipelines further complicate cross-species comparisons.

Sex chromosomes are central to sex determination, and characterizing their evolution provides important insight into changes in sex-determining mechanisms. Available cytogenetic work summarized across Coleoptera indicates that most species possess an XY sex-chromosome system [18], while alternative systems such as X0, complex multi-X systems, and haplodiploidy do occur but are typically phylogenetically restricted to certain groups [18–20]. The X chromosome generally appears to be stable in beetles [14], and is possibly conserved across insects [21]. Evolutionarily old X chromosomes usually take on several unique properties given their sex-biased inheritance and need for dosage compensation to account for Y chromosome degeneration [22, 23]. These properties include evolutionary stability in gene content [24], enrichment for sex-specific or biased genes [25], unique gene expression patterns during meiosis [26], and the formation of unique repeat/TE landscapes. In beetles, it is not known whether the ancestral X has the typical properties of old X chromosomes nor how phylogenetically widespread they may be. [14, 27]. Additionally, fusions between the X chromosome and autosomes have independently generated instances of so-called neo-sex chromosomes in several species of beetle [14–16]. Neo-sex chromosomes have been useful in studying early-stage processes of sex chromosome differentiation and have also been linked to speciation [28–30]. However, most neo-sex chromosomes have been identified through more course-grained cytological approaches, and thus the frequency and phylogenetic distribution of X-autosome fusions across beetles remains unclear.

Here, we take advantage of the rapidly expanding number of publicly available beetle genomes (see Appendix 1 for full list of citations) and we analyze 190 chromosome-level assemblies to (1) broadly characterize genome variation across Coleoptera, (2) evaluate chromosome conservation and the persistence of Stevens elements, (3) identify species with putative neo-sex chromosomes, and (4) explore repeat evolution as a potential driver of karyotypic change. Our analyses reveal that beetles are as genomically diverse as they are species-rich. We observe deep chromosome stability, including conserved synteny and collinearity spanning hundreds of millions of years in some superfamilies, alongside explosive structural variation in others that may be linked to diversification. We further confirm that the X chromosome is broadly conserved across beetles; however, repeated fusions between the X and autosomes, ranging in age and complexity, suggest that neo-sex chromosomes are far more prevalent than previously recognized and may play a substantial role in shaping genome evolution in certain lineages. Together, these results provide a foundation for placing chromosome conservation, genome restructuring, and sex chromosome evolution into a phylogenetic framework across Coleoptera.

## RESULTS

### Beetle evolution and genomic diversity

We analyzed 190 chromosome-level beetle genomes spanning two suborders (Adephaga and Polyphaga), 39 families (16 superfamilies), encompassing more than 300 million years of evolution (Table S1). The final assemblies used in our analyses were all high quality, with most assemblies exceeding 95% BUSCO completeness (Fig. S1; Table S1). These assemblies represent a diverse range of species and span much of the phylogenetic diversity of the order. Karyotypes ranged from four autosomes and XY sex chromosomes, as found in *Endomychus coccineus*, to the more typical nine autosomes and Xyp sex chromosome configuration (small degenerated “parachute” Y) of the model *Tribolium castaneum*, to haplodiploid lineages like *Euwallacea fornicatus* with 39 pseudochromosomes inferred from Hi-C scaffolding (Table S1).

We first explored evolutionary relationships among all taxa by creating individual gene trees (2,124 BUSCOs) and a consensus tree using ASTRAL (Fig. 1A). Phylogenetic relationships among 39 families (16 superfamilies), were found to be generally consistent with previously published large-scale multilocus beetle phylogenies [9, 13]. As expected, given the large number of genes/loci involved in the creation of our consensus tree, bootstrap support was universally high throughout, with 93% of nodes = 1 (Fig. 1A). For superfamily relationships, the only node with support < 1 involved the placement of the Coccinelloidea (bootstrap support = 87); a group with varied placements in other studies [9, 13]. Although bootstrap values were high throughout the tree, these metrics are well known to approach the maximum in phylogenomic datasets, even when the underlying phylogenetic signal is rather weak [31, 32]. Therefore, to further explore the extent of alternative signals in our phylogenetic data, we calculated gene and site concordance factors (gCF and sCF, respectively). These metrics provide a more nuanced view of the underlying genomic signal that contributes to the phylogenetic relationships, in contrast to bootstrap supports (and similar metrics) which provide statistical confidence [32]. Across much of the phylogeny, many branches with high bootstrap support values showed moderate to high levels of discordance (low gCF and sCF) (Fig. 1A). This is not atypical at this evolutionary scale [32], but indicates considerable genomic heterogeneity in certain species rich groups likely due to the rapid radiation over short evolutionary time [12]. Bursts of diversity for certain superfamilies was apparent when exploring internode branch lengths, which often were found to be very short (Fig. S2) suggesting many superfamilies first arose over a short evolutionary timeframe [13, 33]. Thus, the substantial discordance and abundance of alternative gene tree topologies are an important feature of the beetle radiation and will need to be better incorporated in future studies when characterizing trait evolution across this group [32].

**Figure 1.**
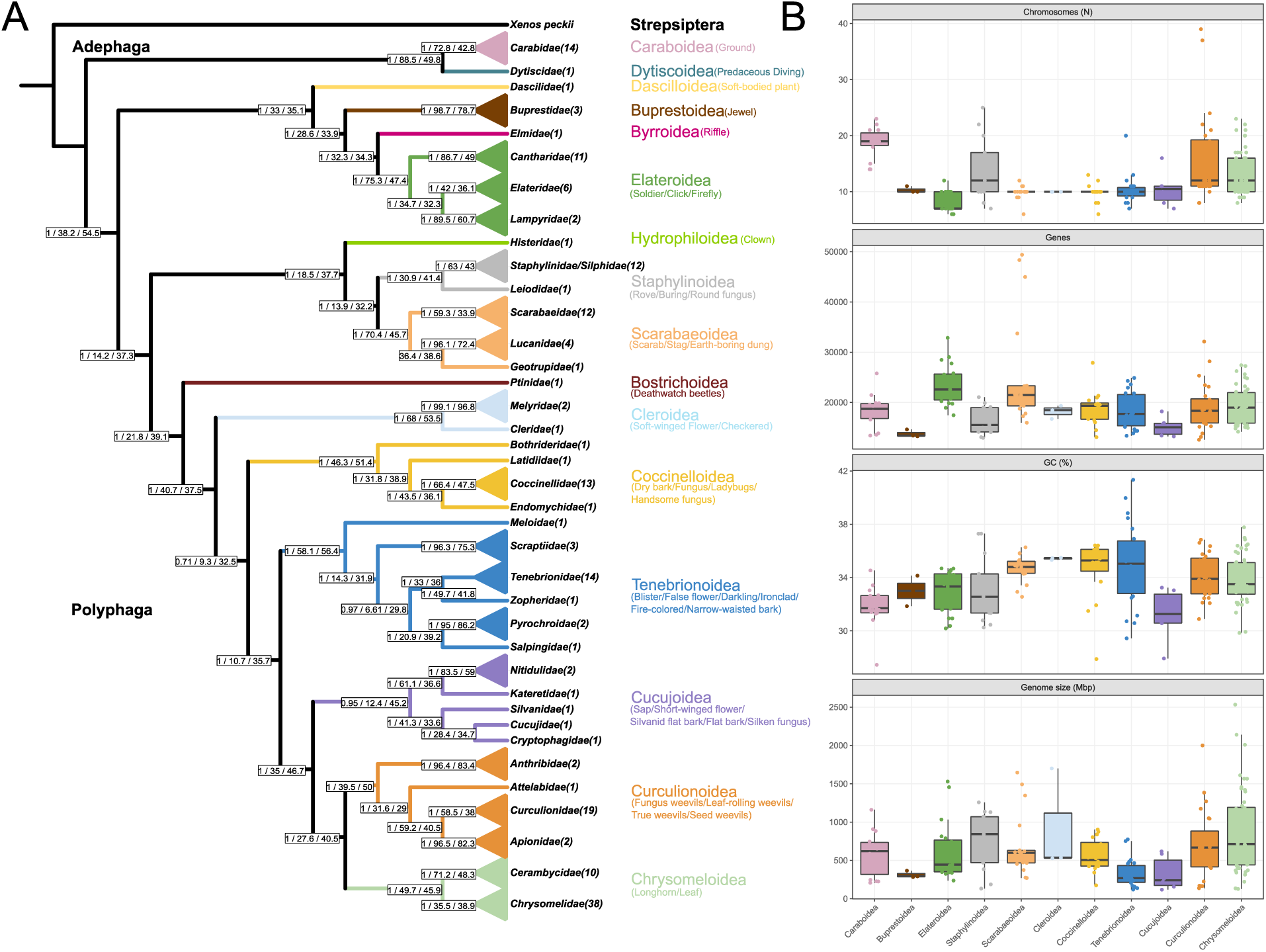
Evolutionary relationships and genome characteristics of beetles. **A)** Phylogenetic relationships of 16 superfamilies (39 families), along with their associated common names (color-coded) inferred from 2,124 BUSCOs. Families represented by multiple species are shown collapsed (triangles), with the number of species in parentheses. Branches show bootstrap support, and gene and site concordance factors, respectively. Note that Staphylinidae and Silphidae were not found to be reciprocally monophyletic, and so were combined and collapsed together. **B)** Genome characteristics for all 190 species, including haploid chromosome number (N), genomic GC%, number of protein coding genes, and assembled genome size (Mb), plotted by superfamily.

We next sought to broadly characterize basic genomic properties such as chromosome number, gene number, assembly length (genome size), and GC content. Some published genome assemblies included high-quality annotations (e.g., *Tribolium castaneum* [34]), but for others, the annotations were either of low quality, were difficult to retrieve, or were altogether unavailable. Therefore, to facilitate our characterizations, and to maximize the set of orthologs for downstream synteny/collinearity analyses, we annotated all genomes using Helixer, a tool that conducts ab initio gene prediction and only requires an unmasked genome as input [35]. We found that the median number of protein coding genes from these annotations across all beetles was 19,039 (range 12,574 to 49,398) with noticeable differences in certain superfamilies, including several outliers in Scarabaeoidea (Fig. 1B). We found haploid chromosome numbers (or pseudochromosomes, see Methods) varied substantially both among and within superfamilies (Fig. 1B). The median chromosome number was 10 (range 5-39) and some lineages have relatively conserved chromosome number, like the Coccinelloidea, Tenebrionoidea, and Scarabaeoidea, while others show pronounced variability like Curculionoidea, Chrysomeloidea, and Staphylinoidea (Fig. 1B). The median genome assembly size when considering only chromosomes from assemblies was 557.981 Mb, but assemblies differed 21.4-fold from the smallest (118.433 Mb) to the largest (2,533.404 Mb) with species in the Chrysomeloidea showing the most variation (Fig. 1B). The GC content was as low as 27.45% to as high as 41.35%, with the Tenebrionoidea showing the most variability (Fig. 1B). In total, our survey of available chromosome-level beetle genome assemblies reveals that not only is Coleoptera a species-rich order, but it is also genomically diverse as well.

### Synteny comparisons with model beetle *Tribolium castaneum* (Stevens elements)

To first explore chromosomal conservation of Stevens elements and to assess chromosome fusions and fissions, we developed a set of complementary visualization and synteny-inference approaches; the first uses a modified implementation of vis_ALG created for identifying conserved Nigon elements in nematodes [36]. Our modified version accommodates beetle assemblies by using BUSCOs assigned to Stevens elements based on *T. castaneum* chromosomal locations. Using our modified vis_ALG, each chromosome (or contig/scaffold) in the reference genome, is plotted with the number of BUSCOs belonging to specific Stevens elements shown in bins based on physical position along the chromosome (or contig/scaffold) (Fig. 2A). This approach allows for straightforward genomic characterization of BUSCO locations and identification of the conserved X chromosome (Fig. 2A).

**Figure 2.**
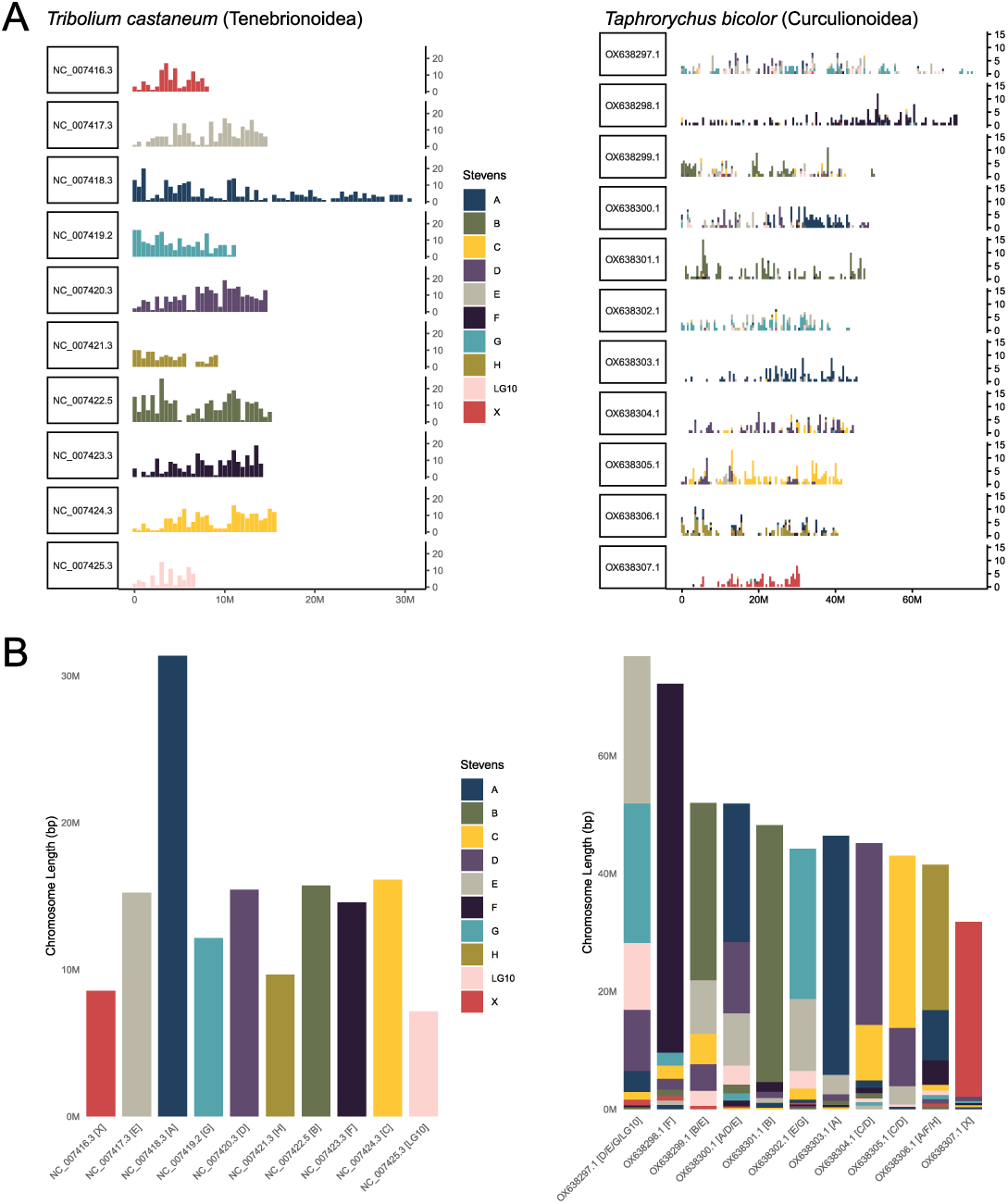
Visualization of chromosome conservation via Stevens elements. **A)** vis_ALG output showing chromosomal location of Stevens element-assigned BUSCOs in *Tribolium castaneum* (reference) and *Taphrorychus bicolor*. Each horizontal panel represents a chromosome in the published reference assembly. **B)** Element_vis output showing the total contribution of each Stevens element to individual chromosomes. The height of each bar corresponds to the physical chromosome length, with colored segments indicating the fraction assigned to each Stevens element. Note that LG10 is a single small chromosome in *T. castaneum* and is not considered a conserved Stevens element.

During initial explorations, we found several species were challenging to interpret, often because of significant changes in genome size, chromosome number, and/or the result of extensive fusions and/or fissions (Fig 2A). To address this, we developed Element_vis, a complementary method that plots each chromosome (or contig/scaffold) by proportion of BUSCOs belonging to particular Stevens elements in descending order (Fig. 2B). This approach allowed for a quick summary of Stevens element proportions for each chromosome and provides an element label if greater than 10% of total BUSCOs on a chromosome are derived from a particular element (Fig 2B). These two tools were designed to summarize chromosome-scale patterns without the need for a full annotation, and one can quickly identify putative Stevens elements and synteny patterns between any beetle and the model *Tribolium castaneum*.

### Identification of putative neo-sex chromosomes

We next set out to identify species with putative neo-sex chromosomes; chromosomes where the conserved ancestral X (Stevens X) appears fused to an autosome or autosomal fragment (Stevens A-H). We first located Stevens element X, which is typically a single, small chromosome (Fig.2), and found all cases where the X chromosome (or pseudochromosome) was also comprised of an additional > 10% of Stevens elements A-H. Species exhibiting these patterns were flagged as possibly having neo-sex chromosomes. This process recovered previously identified neo-sex chromosomes, as well as uncovered novel fusions, including cases of simple X-autosome fusion, as seen in *Dascillus cervinus* (Dascillidae) where the X fused to element F, creating the largest chromosome in the genome (Fig. 3A). Fusions include different autosomal elements, as seen in the stag beetle *Dorcus hopei* (Lucanidae) where the X fused to part of element B (Fig. 3B). While other fusions are quite complex, including the fusion seen in *Ophraella communa* (Chrysomelidae), which suggests a compound chromosome, the result of numerous fusions and fissions of Stevens elements likely prior to the fusion to the X (Fig. 3C). Of the 190 species we explored, we identified 37 (19.5%) species with synteny patterns consistent with neo-sex chromosomes (Table S1, Fig. S3).

**Figure 3.**
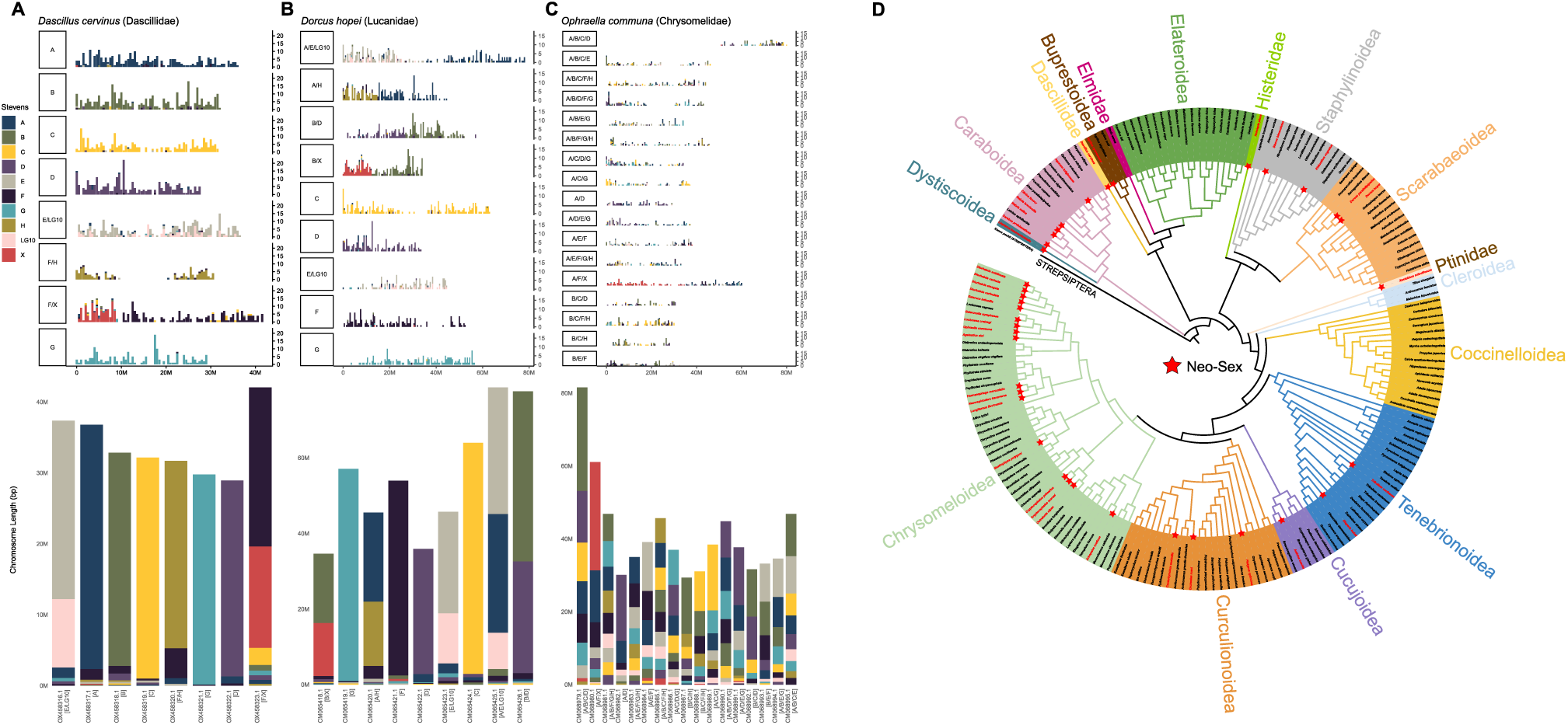
Neo-sex chromosomes in beetles. **A–C** Visualizations of Stevens elements using vis_ALG and Element_vis for three representative beetle species, illustrating distinct X-autosome fusions and increasing complexity in chromosomal organization. **A)** *Dascillus cervinus* with autosomes that correspond to Stevens elements and an X + F neo-sex chromosome. **B)** *Dorcus hopei* with some evidence of Stevens fusion/fission and X + partial B neo-sex chromosome. **C)** *Ophraella communa* with numerous fusions/fissions and X + partial A + partial F neo-sex chromosome. **D)** Phylogenetic distribution of inferred neo-sex chromosomes with colors highlighting beetle superfamilies. Red stars highlight species with putative neo-sex chromosomes identified via vis_ALG and Element_vis.

To better understand the history and distribution of the X-autosomal fusions, we explored fusion patterns in a phylogenetic context (Fig. 3D). Putative neo-sex chromosomes were found in both the suborders Adephaga and Polyphaga, and of the 14 superfamilies we assessed in Polyphaga, 10 had at least one species with a putative neo-sex chromosome. Interestingly, in opposition to the general phylogenetically widespread nature of putative neo-sex chromosomes, the Elateroidea and Coccinelloidea, represented by 19 and 16 genome assemblies, and covering 13 and 15 genera, respectively, did not have a single species with evidence of an X chromosome-autosome fusion. In contrast, the family Chrysomelidae had numerous X chromosome fusions (17 species from 13 genera), although some appear to be the result of ancestral fusion(s) now shared by several species (e.g., species in the genus *Diorhabda*) and therefore likely do not represent independent events. It is important to point out that the species used in these analyses are not from a random sampling of beetle diversity. Many are pests from species-rich families, like Chrysomelidae and Curculionidae, and so certain lineages are overrepresented.

### Broadscale synteny and collinearity in beetles

To complement BUSCO-based Stevens element assignments and provide an independent genome-wide measure of chromosomal conservation, we performed whole-genome synteny (chromosome conservation) and collinearity (gene order conservation) analyses using our genome annotations and GENESPACE [37]. Given the large number of genome assemblies and challenges of characterizing synteny and collinearity over such a large level of divergence, we explored chromosome conservation by performing targeted analyses and contrasts of eight superfamilies (160 species).

We first explored the Tenebrionoidea (Fig. 4A), which contains *T. castaneum*, thereby providing a baseline of chromosomal evolution as one would expect synteny of Stevens elements to be more easily detectable due to lower evolutionary divergence. We found that with only a few exceptions, syntenic blocks were largely maintained across species within the Tenebrionoidea, showing strong conservation of Stevens element structure and consistent chromosome-level correspondence (Fig. 4A). This pattern supports long-term retention of ancestral linkage groups and limited chromosome turnover within the clade, a clade estimated to have formed ∼200 MYA [12, 13] and contains over 33,000 species [8]. We also confirmed the independent fusions that gave rise to neo-sex chromosomes seen in *Tribolium confusum* and *Diaperis boleti* (Fig. 4A).

**Figure 4.**
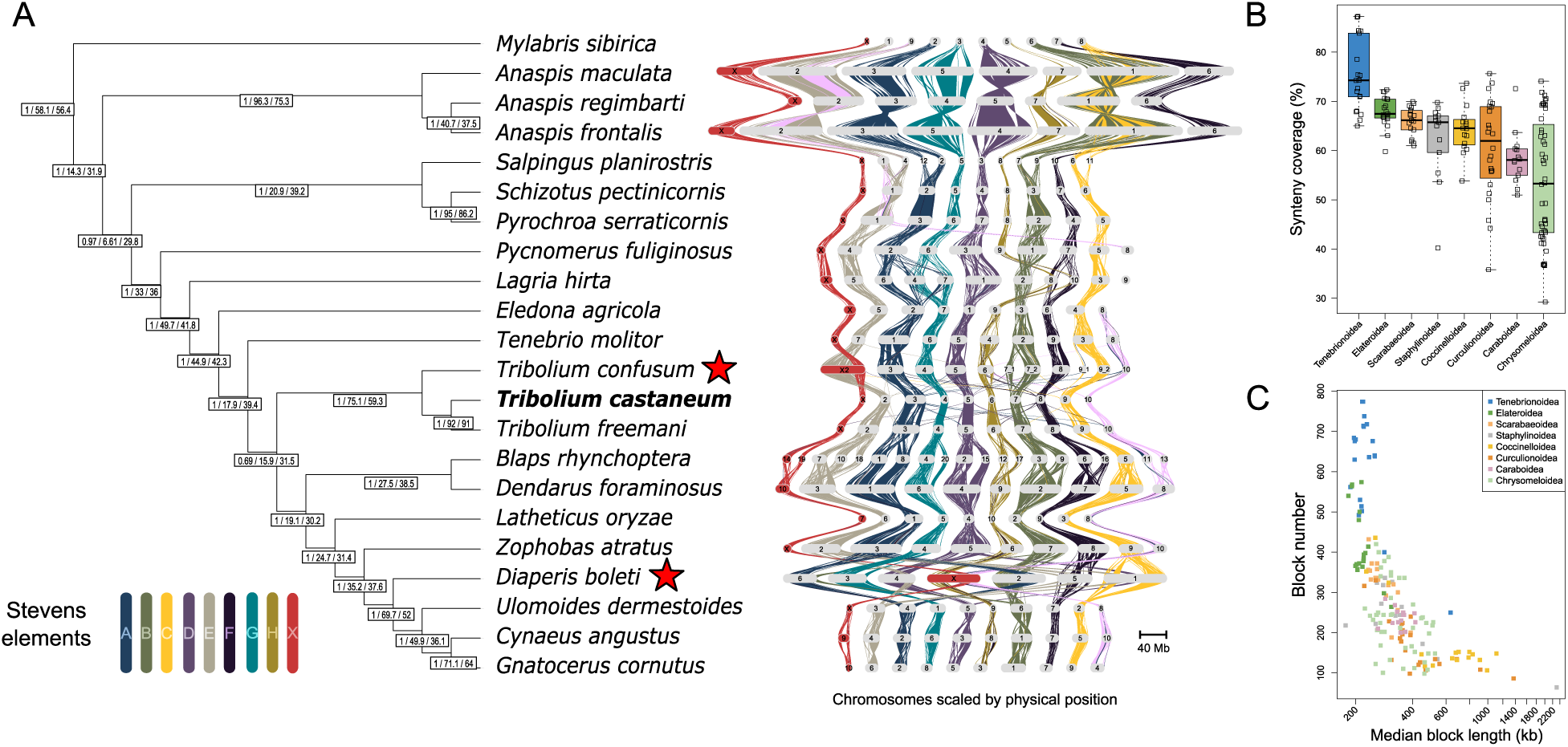
Persistence of Stevens elements in the Tenebrionoidea, yet variable patterns in other superfamilies. **A)** Evolutionary relationships of 22 species in the Tenebrionoidea, with bootstrap support, gene concordance factors (gCF), and site concordance factors (sCF) shown in boxes. Collinearity and synteny across species visualized with a GENESPACE riparian plot color coded by Stevens elements and LG10. Genomes shown scaled by physical position to highlight changes in genome size. Red stars denote X-autosome fusions (neo-sex chromosomes). Note that the *Blaps rhynchoptera* X is likely a single linkage group based on Hi-C [70]. **B)** The percentage of each genome assembly identified as syntenic via GENESPACE in a simple pairwise contrast with *Tribolium castaneum* (Stevens elements) and shown grouped by superfamily. **C)** Relationship between the total number of syntenic blocks identified and median block length in pairwise contrasts with *Tribolium castaneum* (Stevens elements). Species are shown color-coded by superfamily. Note that at this evolutionary scale, no genomes are collinear across a chromosome, and microsynteny increases block number, which generally decreases median block length.

We next sought to quantify the extent to which the conservation of Stevens elements may extend to different beetle superfamilies. We calculated the percentage of each genome that could be identified in syntenic blocks with *T. castaneum* (Fig. 4B). As expected, the percentage of the genome found to be syntenic was high for Tenebrionoidea species (median = 74.3%), and was moderately high (67.5%, 66.2%, 65.7%, and 64.5%) for four other superfamilies: Elateroidea, Scarabaeoidea, Staphylinoidea, and Coccinelloidea respectively (Fig. 4B). Three superfamilies exhibited either lower median values or far more variation, including the Curculionoidea (median = 62%), Caraboidea (median = 58.1%), and Chrysomeloidea (median = 53.3%) (Fig.4B). These superfamily patterns were also reflected in the total number of identified syntenic blocks (Fig. 4C), suggesting superfamilies with low synteny could related to phylogenetic distance (e.g., Caraboidea) or underlying genomic changes (e.g., Chrysomeloidea).

To further assess the conservation of synteny across each superfamily, we jointly interpreted results from GENESPACE riparian plots and modeled pairwise synteny as it relates to the estimated divergence time between species within superfamilies (Fig. S4). We took an approach where synteny analyses were first constrained to Stevens elements (*T. castaneum* reference and used as outgroup) and second where we used a reference from within each superfamily. The purpose of this approach was to allow for the identification of overall trends in chromosome conservation and synteny decay within superfamilies while considering the influence of phylogenetic distance from *T. castaneum* (Tenebrionoidea) on synteny detection. Further, this approach allowed for the characterization of superfamily-specific linkage groups that may differ from Stevens elements.

Results from modeling synteny decay by superfamily are shown in Fig. 5 and Table S2 while corresponding GENESPACE riparian plot images are shown in Fig. 6 and Fig. S5 – S9. In general, across most superfamilies, synteny decreased as pairwise species divergence increased although this pattern was more obvious for some superfamilies (e.g. Elateroidea and Scarabaeoidea, Fig. 5B and C). For the Tenebrionoidea, Elateroidea, Scarabaeoidea, Staphylinoidea and Coccinelloidea, estimates of synteny via Stevens elements were lower than those using superfamily-specific references, but general trends were similar suggesting that phylogenetic distance did decrease overall synteny detection, but not overall rates of synteny decay. Further, in these groups, Stevens elements appear partially to largely conserved based on GENESPACE riparian plots, although in all cases, species with aberrant genomes break trends (Fig 6; Fig S5 – S9).

**Figure 5.**
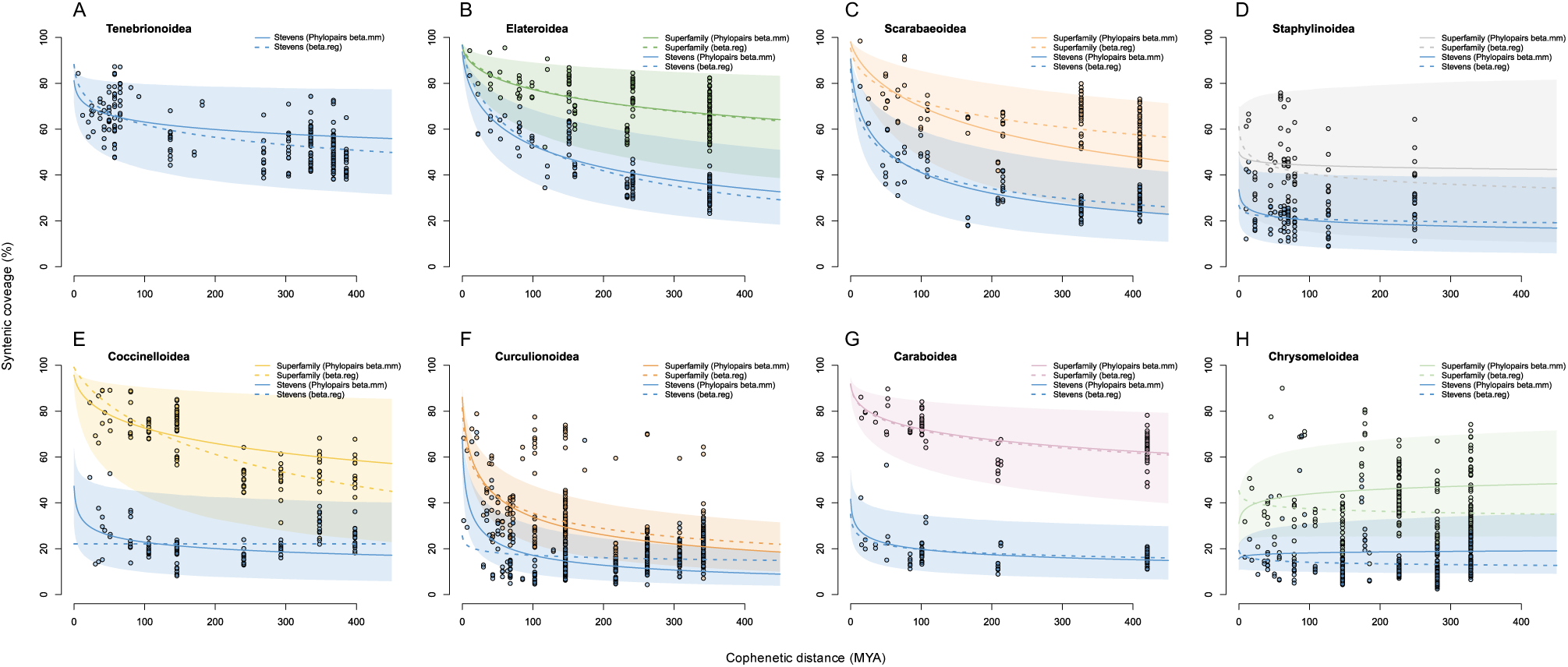
Genome synteny decays over time and varies by superfamily. Relationship between divergence time (cophenetic distance, millions of years ago) and percentage of genome captured in syntenic blocks in species pairs from within eight beetle superfamilies: **A)** Tenebrionoidea, **B)** Elateroidea, **C)** Scarabaeoidea, **D)** Staphylinoidea, **E)** Coccinelloidea, **F)** Curculionoidea, **G)** Caraboidea, and **H)** Chrysomeloidea. Each panel displays beta regression models for two levels of taxonomic comparisons; one where pairwise synteny was constrained to Stevens elements (*T. castaneum* reference; blue) and one where synteny was constrained to a reference from within the superfamily (shown with superfamily specific colors). Solid lines represent phylogenetically-adjusted beta mixture models for lineage-pair traits [66] with dashed lines representing the results from standard beta regressions. Shaded regions represent 95% confidence intervals for the beta mixture models. Full posterior values for both beta models per superfamily can be found in Table S2.

**Figure 6.**
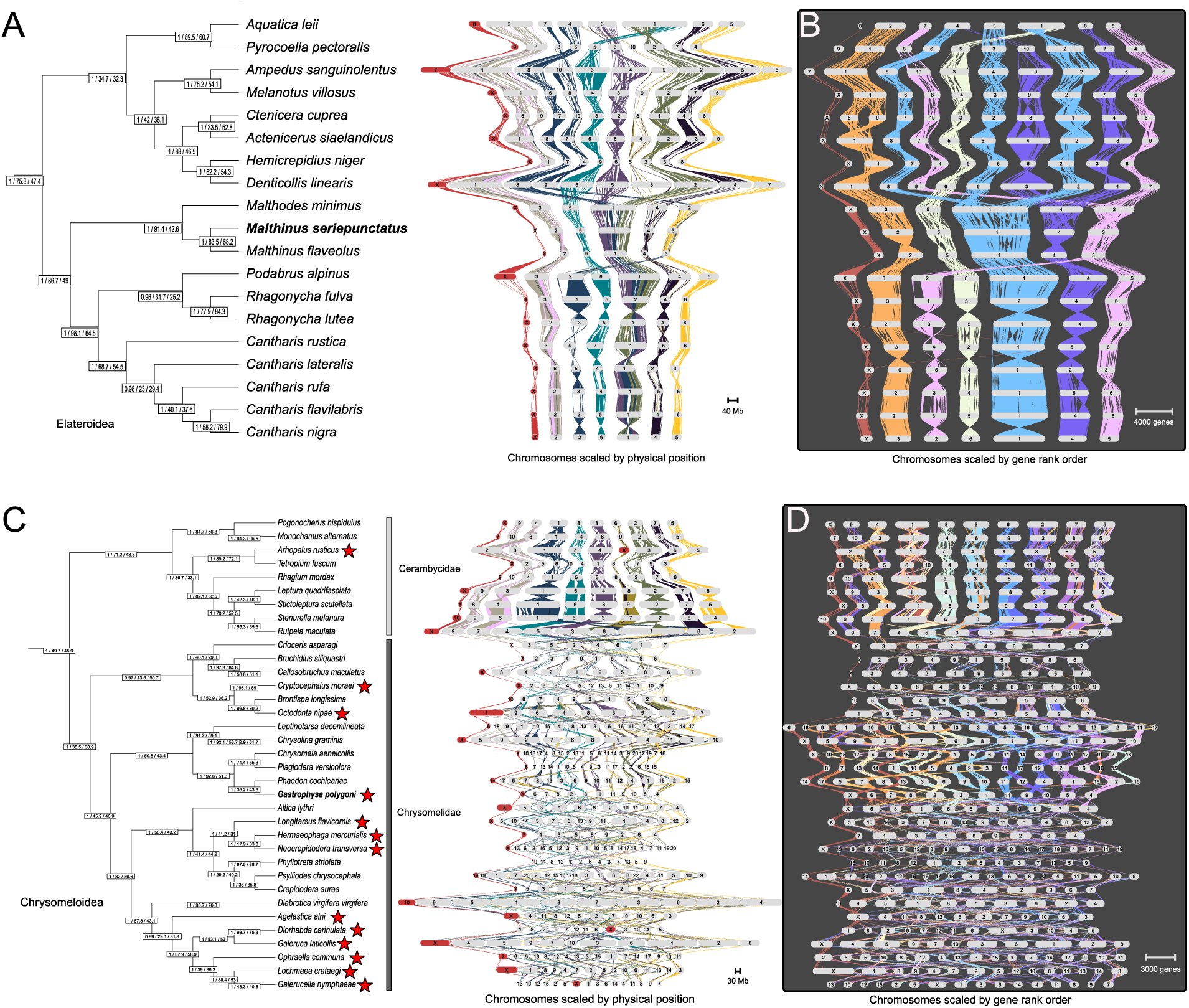
Genome structure evolution trajectories vary across beetle superfamilies. Two groups that exemplify the two extremes seen in beetles are the Elateroidea and Chrysomeloidea. **A)** GENESPACE riparian plot (similar to Fig. 4) for the Elateroidea showing Stevens element conservation and lack of neo-sex chromosomes **B)** Riparian plot using the within-superfamily reference from beta regression analyses (bold, *Malthinus seriepunctatus*) with gene rank order shown to help visualize collinearity. **C)** Riparian plot for the Chrysomeloidea with the Cerambycidae and Chrysomelidae families highlighted to note differences in chromosome conservation and genome evolution between families. **D)** Riparian plot using the within-superfamily reference from beta regression analyses (bold, *Gastrophysa polygoni*) with gene rank order shown to help visualize collinearity. Note the *Arhopalus rusticus* genome shown here corresponds to an earlier assembly version, which incorrectly labeled the X. Putative neo-sex chromosomes highlighted with red stars.

Our analyses revealed three superfamilies of note, each for different reasons: the Curculionoidea, Caraboidea, and Chrysomeloidea. In Curculionoidea, synteny appears to quickly decay between species pairs (Fig. 5F) and synteny was challenging to recover for many species using either Stevens elements or the superfamily-specific reference approach (Fig. S8). In Caraboidea, although some linkage groups were consistent with Stevens elements, estimating synteny using a within superfamily-reference greatly improved synteny contrasts suggesting that the deep Adephaga-Polyphaga split (∼300 mya) might be the phylogenetic limit of Stevens element conservation (Fig. 5G and Fig. S9). For the Chrysomeloidea, low, and highly variable levels of synteny between species pairs were found at all levels of divergence (Fig. 5H) and clear patterns of synteny between many species were challenging to recover.

No contrast better exemplifies the range of variation seen across beetles than the differences between the Elateroidea and the Chrysomeloidea (Fig 6). The Elateroidea broadly exhibits conserved Stevens elements, with only a few simple chromosomal fusions or fissions within certain lineages (Fig. 6A). Finer-scale analyses using the within-superfamily reference reveal numerous species-specific inversions, yet overall chromosomes appear largely conserved (Fig. 6B). In contrast, the Chrysomeloidea, and specifically the family Chrysomelidae, demonstrate a clear loss of Stevens elements and drastic changes in genome structure (Fig. 6C, Fig. S10A). Within the Chrysomelidae, chromosome numbers and genome sizes were found to be highly variable, with numerous fusions and fissions driving the redistribution of syntenic blocks across chromosomes including numerous neo-sex chromosomes. Changes in collinearity across chromosomes and species appear to be so substantial that analyses using the within-superfamily reference also struggled to connect syntenic regions between many species in this group (Fig. 6D). Interestingly, the sister family, Cerambycidae appears to largely retain Stevens elements and shows much slower synteny decay over time (Fig. 6C, Fig. S10B).

### Repetitive elements as drivers of chromosomal evolution?

Repetitive elements, and specifically transposable elements (TEs), are known to contribute to overall genome size and have also been implicated in facilitating structural changes like inversions [38] and large-scale karyotypic changes [39]. We set out to use a universal pipeline, Earl Grey [40], to identify and characterize the repeat landscape from a subset of beetle species to better understand repeat dynamics and potential links to chromosomal fusion/fission and karyotypic change. We selected 71 species in total, representing the full phylogenetic breadth of our dataset and using species with a wide range of genome sizes and karyotypic configurations (Fig. 7A, Table S1). We found the proportion of the genome masked for repeats was as low as 13.0% (*Nicrophorus investigator*) to as high as 78.5% (*Tillus elongatus*) (Fig. 7A). The relative proportions of common repeats differed substantially across species, with no obvious phylogenetic patterns associated with chromosome stability and karyotype evolution (Fig. 7A). Of identifiable TEs, DNA transposons typically make up the largest fraction (mean = 23.1%, SD= 11.1%), followed by LINE elements (mean = 15.2%, SD = 11.1%) (Fig. 7A). However, significant variation occurs across species and among all superfamilies (Fig. 7A). LTR elements (mean = 8.3%, SD = 5.8%) and simple repeats (satellites; mean = 7.7%, SD = 9.3%) appear quite variable in beetles, with little clear phylogenetic pattern. We found a weak, but positive relationship between total genome size and DNA transposons and LINE elements and a negative relationship with simple repeats (Fig. S11). We found no relationship between LTR elements and genome size (Fig. S11). Thus, on this broad phylogenetic scale, repeat evolution appears highly variable and rather idiosyncratic with few clear trends. It is important to point out that many repeats remain unclassified in beetles and additional research is needed to further characterize the full beetle repeat-ome and its evolution.

**Figure 7.**
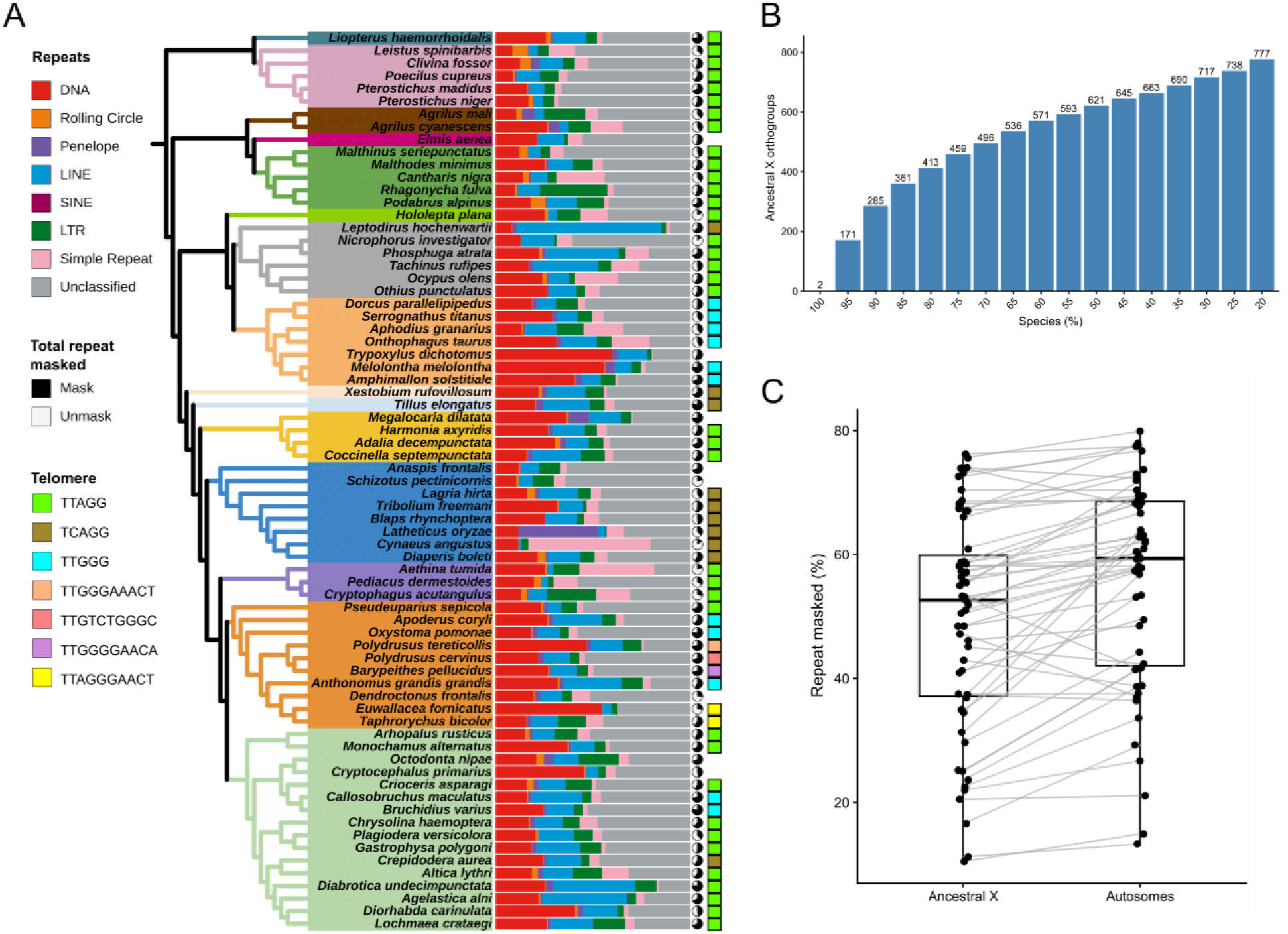
Repeat evolution and ancestral X chromosome stability. **A)** Repeat landscape for 71 species showing the proportional contribution of major repeats (DNA, Rolling Circle, Penelope, Line, Sine, LTR, Simple Repeat, and Unclassified). Pie charts show the proportion of the genome assembly that was repeat masked. Putative telomere sequence motifs are represented by colored squares. **B) N**umber of shared X-linked genes using a range of thresholds (20-100%). Each threshold represents the minimum proportion of species in which an orthogroup must be X-linked to be considered conserved. **C)** Paired boxplot comparing the percentage of bases masked for transposable elements between the ancestral X chromosome and autosomes from the same species.

Previous studies of telomeric sequence in beetles suggest several motifs are present broadly across the radiation, with (TTAGG)_n_ likely being the ancestral sequence in insects and found in some beetles, but with several transitions to alternatives in various superfamilies [41]. To find known telomere repeats, identify new telomeric motifs, and explore phylogenetic patterns associated with diversification and genome evolution, we also ran tidk [42] on each genome assembly (n = 190). We were able to confidently identify the putative telomeric repeat in 67% (n = 128) of the assemblies (Table S1). We found most genomes contained one of three core telomeric motifs: 67% of species harbored the putative ancestral motif of insects (TTAGG)_n_, while 13% were (TCAGG)_n_, and 13% (TTGGG)_n_. Several other longer motifs that contained these three strings, or were quite similar in sequence, were present in lower frequencies and were restricted to the Curculionoidea (Table S1). Phylogenetic conservation of certain motifs was evident, and several superfamilies contained only one motif (e.g., Tenebrionoidea/TCAGG: n = 13, Caraboidea/TTAGG: n = 11, Elateroidea/TTAGG: n = 15, Scarabaeoidea/TTGGG: n = 11), while several others had some variability yet still a dominant motif (e.g., Chrysomeloidea, 91% = TTAGG) (Fig. 7A, Table S1). The only obvious standout was Curculionoidea which uses several unique long motifs in addition to two of the three core motifs (Fig. 7A, Table S1). In total, our results show that across beetles, there is significant telomeric sequence diversity and transitions among closely related superfamilies. However, we found no clear association between telomere diversity or turnover consistent with patterns of karyotype evolution as highlighted by superfamilies like Chrysomeloidea (Fig. 7A, Table S1).

### The beetle X chromosome

Our results show that the ancestral (unfused) X has been conserved for over 300 million years in beetles (and probably longer across insects [21]) and has fused numerous times to autosomes. We find that the ancestral X of beetles on average harbors ∼1k genes, which typically accounts for ∼5% of all genes in the average genome (Fig. S12). When fusions between the ancestral X and autosomes occur, this results in a gain in gene content, on average doubling the proportion of the genome that is sex-linked (Fig. S12). The long-term conservation of the ancestral X in beetles is reflected by its stable gene content, and we have identified core genes shared across species depending on our filtering stringency (Fig. 7B). For example, 90% of species examined share 285 orthologous genes on their X chromosomes, when allowing for X gene duplications. A slightly relaxed threshold (70% of species) identifies nearly 500 genes shared on the X (Fig. 7B). Thus, the long-term stability of gene content suggests that many genes have been evolving on an X for hundreds of millions of years, if not even longer [21]. Interestingly, in addition to gene stability, we find that TE concentration typically differs between the ancestral X and autosomes. For most species (84%, n = 46 of 55), the X has fewer bases masked for TEs when compared to the autosomes (Fig. 7C), a pattern noted previously but from only a few species [14, 43, 44]. This difference appears to be a persistent characteristic of the beetle X, one that has withstood repeated changes in overall genome size and TE family abundance during diversification into the ∼400k described extant species (Fig. 7A). Thus, with respect to gene and repeat content, the ancestral X stands out as distinct within the context of a beetle genome.

## DISCUSSION

Beetles have existed for roughly 300 million years and are extraordinarily diverse, and this species richness is matched by substantial genomic diversity. By interrogating 190 chromosome-level genome assemblies, standardizing annotations, and applying comparative analyses in a phylogenetic framework, we modeled how synteny decays through time and assessed genome evolution and chromosome conservation across one of the most successful animal radiations on Earth. We find that beetle genomes vary widely in size, chromosome number, GC content, telomere motifs, and the composition and density of transposable elements.

A central goal of this study was to determine the extent of chromosome conservation in beetles and whether Stevens elements are maintained across the radiation, analogous to Muller elements in flies (Diptera) [6] and Merian elements in butterflies and moths (Lepidoptera) [7]. Our analyses reveal a nuanced picture of beetle genome evolution: deep conservation of ancestral linkage groups in some large, old superfamilies (e.g., Elateroidea and Tenebrionoidea) contrasted with repeated structural change within a family (e.g., Chrysomelidae). Across most of Polyphaga, ancestral linkage groups remain largely stable, and the genomes of most species can be explained by simple fusions and fissions among Stevens elements. However, our results suggest that chromosome-scale synteny may erode faster in beetles than in Lepidoptera, where Merian elements appear more stable [7, 45]. In addition, gene order (collinearity) appears to change rapidly in beetles, with inversions frequently separating closely related species or genera, similar to patterns reported in *Drosophila* [46, 47]. Together, these results add to growing evidence that many animal groups experience long-term constraints on chromosome number and linkage-group structure, even though some lineages occasionally deviate markedly [1, 4].

What allows some lineages to escape these apparent constraints and diverge rapidly in genome structure remains unclear. A productive way to address this question is to compare closely related clades that differ strongly in chromosome conservation. Prior comparisons of three chrysomelid genomes to other beetle families suggested that some leaf beetles lack broad chromosome-level conservation [48]. Our results show that this pattern extends broadly across Chrysomelidae, but appears largely absent in Cerambycidae. These sister families diverged ∼150 MYA and have comparable extant species richness (Chrysomelidae ∼32,500 species; Cerambycidae ∼30,000 species). We suggest that targeted comparisons of the genomic features that distinguish these lineages could be particularly informative for identifying drivers of fusion/fission frequency and broader differences in genome evolution.

Beyond illuminating genome evolution, placing beetle chromosomes into a Stevens-element framework has practical benefits for comparative genomics because *T. castaneum* remains the primary functional model for Coleoptera, and much functional annotation is anchored to this species [49, 50]. A shared naming scheme can help map genes and chromosomes across species in an explicit evolutionary context, much as Muller elements do in flies to facilitate studies of genome evolution and sex-chromosome turnover [47, 51, 52]. At the same time, our results highlight limits to this approach: in some groups (e.g., Curculionoidea and Chrysomeloidea), extensive rearrangement and reduced detectable synteny make Stevens-element assignments less straightforward and, in some cases, less informative.

Any association between repeat evolution and large-scale karyotypic change across superfamilies was not obvious, and it may be difficult to detect at the evolutionary scale of our analysis. Repeat content varies substantially across beetles, although the broad trend that larger genomes tend to harbor more repetitive DNA is consistent with patterns in other insects [53]. Our telomere analyses are broadly consistent with recent studies documenting extensive turnover of telomeric motifs across Coleoptera [54], but how telomere evolution could relate to large-scale chromosomal change remains unclear. Beyond the general pattern that DNA transposons are the most abundant TE class in many species, followed by LINEs [55], few consistent trends emerged across superfamilies. A notable challenge is the high fraction of unclassified repeats in many assemblies, which in some species accounts for most masked sequence. This is particularly evident in Adephaga, where several species are >50% repeat-masked and dominated by unclassified repeats. Further work to identify and classify beetle TEs is therefore an important prerequisite for testing mechanistic links between repeat dynamics and chromosomal evolution across Coleoptera [55].

The proportion of species in our dataset with patterns consistent with neo-sex chromosomes (∼19%) is substantially higher than estimates from cytological karyotyping, which suggest neo-XY systems account for ∼5% of beetle karyotypes (n = 209 of 4,322 records with sex chromosome system reported) [18]. Several factors could contribute to this difference, most obviously differences in taxon sampling between datasets. In addition, our inference is based on assembly-derived linkage patterns relative to the conserved X, and most candidates lack independent validation (e.g., male vs. female coverage tests) to confirm sex linkage [56]. Nonetheless, there are reasons to suspect that cytology-based estimates may underestimate the true frequency of X–autosome fusions. In particular, older fusions in which the neo-Y has largely degenerated, fusions involving small chromosomes, or fusions to small autosomal fragments can be difficult to detect in chromosome squashes. Consistent with this, assembly-based analyses in Lepidoptera have also uncovered many neo-sex chromosomes that were previously unrecognized [7].

Of the 37 species we identified with putative neo-sex chromosomes, only 11 are represented in the Coleoptera karyotype database [18], and only three were previously reported as harboring neo-sex chromosomes (*Diaperis boleti*, *Dorcus parallelipipedus*, *Tribolium confusum*). The remaining eight (*Agonum fuliginosum*, *Carabus problematicus*, *Clivina fossor*, *Cryptocephalus moraei*, *Cryptocephalus primarius*, *Nebria brevicollis*, *Neocrepidodera transversa*, *Ophraella communa*) are listed as XY, XO, or Xyp in the database. Four of these species are carabids (Caraboidea; Adephaga), a deeply divergent clade for which our synteny and collinearity signals were comparatively variable; these candidates therefore warrant additional scrutiny and independent confirmation. Even with these caveats, our results suggest that neo-sex chromosomes may be more common in beetles than previously appreciated and that cytological data alone may substantially underestimate the frequency of sex chromosome–autosome fusions.

The distribution of putative neo-sex chromosomes does not appear random across the beetle radiation. Instead, candidates cluster in superfamilies with evidence of rapid chromosomal evolution, such as Chrysomeloidea (especially Chrysomelidae) and Curculionoidea, suggesting that lineage-specific differences in overall turnover rate may predispose some clades to repeated sex chromosome–autosome fusions. Recent work in scarab beetles further suggests that such fusions can occur more frequently than expected by chance, potentially because they help resolve sexually antagonistic selection [57]. Future studies that combine assembly-based identification of neo-sex chromosomes with independent validation and phylogenetic models of transition rates should clarify both how often these fusions arise and what evolutionary forces favor them in particular lineages.

## METHODS

### Genome filtering

Beetle genome assemblies identified as “chromosome-level”, were retrieved from the National Center for Biotechnology Information (NCBI) genome database on October 29, 2024. Metadata such as species name, accession number, genome size, chromosome number, karyotype information, assembly level, and taxonomic classification were initially extracted from NCBI and stored for downstream filtering and analysis (Table S1). Of the initial 218 assemblies downloaded, only 200 were retained as several species had multiple assemblies. Prior to all downstream analyses, we filtered out any unplaced contigs, scaffolds, mtDNA, and Y chromosomes.

Genome assembly quality was assessed using Benchmarking Universal Single-Copy Orthologs (BUSCO v5.7.1) with the *endopterygota_odb10* lineage dataset. During initial BUSCO analyses, we had issues analyzing nine assemblies, and they were excluded from further analysis due to unrecognized FASTA headers or issues with contigs that prevented analysis. One additional genome (*Pogonas chalceus*) was further removed due to exceptionally low completeness, with nearly 50% of BUSCOs missing. The final dataset comprised 190 chromosome-level beetle genomes used in all subsequent analyses (Table S1).

### Phylogenetic tree

To infer evolutionary relationships among beetle species, we used the BUSCO_phylogenomics pipeline available at https://github.com/jamiemcg/BUSCO_phylogenomics. This pipeline extracted all single-copy orthologs identified by BUSCO present in at least four species, performed multiple sequence alignment using MUSCLEv5.1 [58], and trimmed low-quality regions with trimAlv1.5.1 [59]. Gene trees were inferred for each ortholog set using IQ-TREE3v3.0.1(https://github.com/iqtree/iqtree3). The resulting collection of gene trees were then combined and used as input for ASTRALv5.7.8 [60] to generate a species consensus tree with *Xenos peckii* (Strepsiptera) used as the root. Tree robustness and congruence were evaluated through bootstrap support, gene concordance factors, and site concordance factor likelihood estimates using IQ-TREE3 and visualized using Interactive Tree of Life (iTOL v7) [61] .

### Independent annotation and characterization of beetle genomes

Each beetle genome assembly was independently annotated using Helixer [35], a deep learning–based eukaryotic genome annotation tool that integrates neural network predictions with hidden Markov models to identify gene models from unmasked genomic sequence data. Annotations were generated under default parameters with their invertebrate lineage model (invertebrate_v0.3_m_0100.h5). Genome completeness was assessed using BUSCO [62]. From each annotated genome, genome size, gene count, and GC content were calculated using R and Python scripts designed to parse Helixer outputs and corresponding FASTA files. The haploid chromosome number for each species was determined from the NCBI genome assemblies and from exploring the associated genome notes when available (see Table S1). Karyotype information was determined using the Coleoptera Karyotype Database [18]. If no karyotype was reported or available for a specific species, then the haploid chromosome number represents a pseudochromosome number reported in the genome note and on NCBI. Where the haploid chromosome (or pseudochromosome) number and the reported karyotype were inconsistent (n = 7), the pseudochromosome number was used, and the discrepancy noted in Table S1.

### Pairwise contrasts with *Tribolium castaneum* (Stevens elements)

To characterize chromosomal conservation of Stevens elements across beetles, we modified the R package vis_ALG (https://github.com/pgonzale60/vis_ALG), originally developed to visualize Nigon elements [36], to accommodate beetle (and fly) genome assemblies. BUSCO genes were assigned to Stevens elements based on their location in *Tribolium castaneum*. The chromosomal position of each BUSCO was used to infer the Stevens element ortholog in each species’ assembly. Each chromosome that was at least 500kb, are divided into bins of equal physical length and the count per Stevens element was calculated within each bin. An additional script (element_vis.R) plots chromosomes, using the same length filter, vertically and shows the proportion of BUSCOs derived from each Stevens element in descending order. Chromosomes were then labeled with which Stevens elements comprised more than 10% of total BUSCOs on that chromosome.

### Identification of putative neo-sex chromosomes

For each genome assembly, we used our two scripts (described above) to first identify Stevens X, which is typically a distinct single linkage group. We then looked for cases where the X chromosome contained genes from autosomal Stevens elements (A–H). Chromosomes were classified as putative neo-sex chromosomes when more than 10% of BUSCOs assigned to that chromosome were derived from one or more autosomal Stevens elements. Species exhibiting these patterns were flagged as candidates for further inspection in downstream analyses.

### Multispecies synteny and collinearity comparisons

To investigate patterns of chromosomal conservation and structural rearrangement across Coleoptera, we performed multispecies synteny and collinearity analyses using GENESPACEv1.3.1 [37] and OrthoFinderv2.5.5 [63]. Prior to analysis, duplicated congeners were removed (N = 13) within Chrysomeloidea to reduce redundancy. For each genome, protein sequences from our Helixer annotations were used as input for OrthoFinder to infer orthogroups, which were subsequently used by GENESPACE to identify collinear gene blocks across genomes within each superfamily.

### Quantifying syntenic coverage and its relationship to divergence time

To quantify synteny, collinearity, and the conservation of Stevens elements, we parsed GENESPACE output files corresponding to phased syntenic blocks (i.e., *_phasedBlks.csv); files that contain genomic coordinates for all inferred syntenic blocks between pairs of genomes. Two complementary datasets were analyzed for each superfamily: (1) a dataset including *Tribolium castaneum* (Tribolium_castaneum_GCF_000002335.3) as a reference genome and outgroup, (2) a dataset excluding *T. castaneum*, in which a representative species within the superfamily was used as the reference. In effect, approach 1) constrained synteny analyses to genomic blocks that were able to be found in *T. castaneum* while 2) used a more closely related within-family species to explore synteny in the event that evolutionarily distant contrasts of the former approach failed. For Tenebrionoidea, only within superfamily comparisons were performed as it contains *T. castaneum*.

To ensure consistent measurement of syntenic block size independent of orientation, genomic coordinates for each block were normalized by defining start and end positions as the minimum and maximum of the reported coordinates. GENESPACE can report fragmented or overlapping intervals within a single syntenic block (duplicated blkID), this could lead to double counting when estimating block lengths. To address this, we merged all overlapping intervals within each chromosome of a given block before summing across the chromosomes, producing an estimate of total genomic block length. For each pairwise genome comparison, we quantified: (1) the total number of syntenic blocks, (2) the distribution of block lengths, and (3) the total genomic coverage of syntenic regions. Genomic coverage was calculated as the total non-overlapping base pairs contained within syntenic blocks for each genome. To enable comparisons across species with varying genome sizes, syntenic coverage was normalized by genome size, resulting in the proportion of each genome contained within collinear blocks.

### Syntenic patterns using beta-mixed model regression

To relate synteny patterns between species to evolutionary divergence, we converted our tree to an ultrametric time-calibrated species tree using the chronos function in the R package ape v5.8.1 [64]. Divergence times for Coleoptera suborders (Adephaga and Polyphaga) and infraorders (Elateriformia, Staphyliniformia, Cucujiformia) were taken from Table 1 in [13]. Divergence times for families (Carabidae, Scarabaeidae, Cerambycidae, Chrysomelidae, Curculionidae, Coccinellidae, Tenebrionidae, and Cantharidae) were extracted from [65] time trees using the age range across all four chains (IR chain 1, IR chain 2, AC chain 1, and AC chain 2) as calibration windows. Pairwise phylogenetic distances were then calculated from this tree using the cophenetic.phylo function in ape, where each pairwise distance represents twice the age of the most recent common ancestor (2 × MRCA) in millions of years. Distances were log-transformed as log(d+1) prior to modeling to account for the right-skewed distribution resulting from uneven lineage sampling across the radiation. Syntenic coverage percentages were rescaled to open unit interval using the method described in [65]:

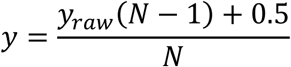

*N*where N is the number of pairwise observations. To account for the non-independent pairwise comparisons arising from shared evolutionary history, a lineage-pair covariance matrix (C_M_) was constructed using the taxapair.vcv function in phylopairsv0.1.1 [66] under the default square difference model. However, due to the scale of the covariance matrix model, “σ^2^_scale” was not identifiable without constraining the matrix [67]. To do this the covariance matrix was trace-normalized to represent the proportion of variance explained by each component by dividing the entire matrix by the sum of the diagonal elements:

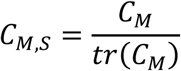

### Model Fitting and Inference

Two beta regression models were fit to each dataset using the betareg.stan function in phylopairs. The first is a mixed model (beta.mm) with the default logit link function and using the trace-normalized C_M_. The second is a standard beta regression (beta.reg) with a logit link function and no phylogenetic correction. This allows us to have a direct comparison of slope estimates with and without the covariance correction. Each model was run twice per superfamily, once using *T. castaneum* as the reference genome to represent Stevens element conservation across the superfamily, and once using within-superfamily reference genomes to characterize synteny decay over evolutionary time. Posterior inference was performed using 10,000 iterations across four chains with adapt_delta = 0.99 and max_treedepth = 15. Fitted values and 95% credible intervals of each superfamily were visualized by generating predictions across 450 million years of cophenetic distance. The prediction values were back transformed from logit scale to percentages producing a fitted curve. All posterior values associated with either beta.mm or beta.reg are reported in Table S1.

### Repetitive sequence identification

To find and characterize repetitive sequence in each genome, we used the Earl Grey v7.0.1 pipeline [40] with default settings. Earl Grey is an automated TE annotation pipeline built largely on output produced from RepeatModeler2 [68] and RepeatMasker [69] and produces numerous output files by default. To extract the relative proportions of TE classifications found within each genome assembly, we parsed the default summary highLevelCount text file and extracted the number of bases annotated for major repeat categories DNA, Rolling Circle, Penelope, LINEs, SINEs, LTR, simple repeats, and Unclassified repeats. Additionally, we used the same file to parse the total percentage of the genome that was masked.

To identify telomeric sequences, we ran each genome assembly through tidk v0.2.65 [42] using default settings. We plotted the frequency of common beetle telomere motifs of all 190 genomes using a script that processes output files and shows forward, reverse, and total telomeric repeat counts across 10kb genomic windows. The criteria used to determine the telomeric repeat motif were: if the motif had greater than 100 copies, was on at least one end of the chromosome, and appeared on at least 3 chromosomes. We then manually inspected the flagged chromosomes using NCBI and confirmed the repeat motif and extended the telomeric sequence when the flagged motif was a substring of a larger repeat as suggested in [42]. Results for all species are reported in Table S1. Visualization of phylogenetic patterns of repeats (TEs and telomeric sequences) was done using iTOL [61].

### Beetle X chromosome gene content and repetitiveness

To assess conservation of X-linked gene content, we used *Tribolium castaneum* as the ancestral X reference and identified orthologs across assemblies using OrthoFinder. We evaluated how consistently each gene remained X-linked across 153 species (excluding 37 neo-sex species) using a range of minimum-proportion thresholds (20-100%), where each threshold represents the minimum proportion of species in which an orthogroup must be X-linked to be considered conserved. In order to compare repeat content between chromosomes, we extracted transposable element annotations from Earl Grey output files associated with the ancestral X, excluding satellite repeats. This process was repeated for autosomes. The percentage of bases masked for transposable elements between the ancestral X chromosome and autosomes for each species was compared using paired boxplots showing per species relationship.

## DATA AVAILABILITY

All genome annotations and output needed to recreate our analyses are available at Figshare (https://figshare.com/projects/Datasets_for_Species-rich_and_genomically_diverse_comparative_genomics_reveal_how_fusions_fissions_and_sex_chromosomes_have_shaped_beetle_evolution_/274777). Relevant code is available at https://github.com/DwayneTally/Stevens-element-analysis https://github.com/DwayneTally/Elements_vis_ALG

## ACKNOWLEDGEMENTS

We thank the numerous genome sequencing projects that have made these genomes assemblies available to the public that form the foundation of our research. Two need special mention including the Tree of Life Programme (https://wellcomeopenresearch.org/treeoflife) and the USDA I5K project https://i5k.nal.usda.gov/. We also thank M. Hahn for suggestions on several analyses and G. Lagunas-Robles, G. Coffing, A. Preble and T. Johnson for comments on earlier versions of this manuscript. This work would have been impossible without help from the Indiana University UITS high-performance computing cluster, which is supported in part by Lilly Endowment, Inc., through its support for the Indiana University Pervasive Technology Institute. This research was supported by NIH NIGMS MIRA grant: R35GM151123.

## Notes

### Competing Interest Statement

The authors have declared no competing interest.

