## Appendix for "Species-rich and genomically diverse: comparative genomics reveals how fusions, fissions, and sex chromosomes have shaped beetle evolution"

**Genome assembly references:**


Ang, G., Zhang, A., Obrycki, J., & Sethuraman, A. (2024). A high-quality genome of the convergent lady beetle, Hippodamia convergens. *G3 (Bethesda)*, *14*(6). https://doi.org/10.1093/g3journal/jkae083

Arnqvist, G., Westerberg, I., Galbraith, J., Sayadi, A., Scofield, D. G., Olsen, R. A., Immonen, E., Bonath, F., Ewels, P., & Suh, A. (2024). A chromosome-level assembly of the seed beetle Callosobruchus maculatus genome with annotation of its repetitive elements. *G3 (Bethesda)*, *14*(2). https://doi.org/10.1093/g3journal/jkad266

Ashworth, M., Darwin Tree of Life Barcoding, c., Wellcome Sanger Institute Tree of Life, p., Wellcome Sanger Institute Scientific Operations, D. N. A. P. c., Tree of Life Core Informatics, c., & Darwin Tree of Life, C. (2023). The genome sequence of a cockchafer, Melolontha melolontha (Linnaeus, 1758). *Wellcome Open Res*, *8*, 222. https://doi.org/10.12688/wellcomeopenres.19434.1

Barclay, M., Gurney, M., Natural History Museum Genome Acquisition Lab, Darwin Tree of Life Barcoding Collective, Wellcome Sanger Institute Tree of Life Management, S. a. L., team, Wellcome Sanger Institute Scientific Operations: Sequencing Operations, Wellcome Sanger Institute Tree of Life Core Informatics team, & Tree of Life Core Informatics collective, D. T. o. L. C. (2026). The genome sequence of a weevil, Sitona lineatus (Linnaeus, 1758). *Wellcome Open Res*. https://doi.org/10.12688/wellcomeopenres.25280.1

Barclay, M. V. L., Crowley, L. M., McCulloch, J., & Telnov, D. (2025). The genome sequence of the two-banded mould beetle, Cartodere (Aridius) bifasciata (Reitter, 1877). *Wellcome Open Res*. https://doi.org/10.12688/wellcomeopenres.23743.1

Barclay, M. V. L., Geiser, M., Matsumoto, K., Pash, E., Natural History Museum Genome Acquisition, L., Darwin Tree of Life Barcoding, c., Wellcome Sanger Institute Tree of Life Management, S., Laboratory, t., Wellcome Sanger Institute Scientific Operations: Sequencing, O., Wellcome Sanger Institute Tree of Life Core Informatics, t., Tree of Life Core Informatics, c., & Darwin Tree of Life, C. (2024). The genome sequence of the Judas Tree Seed Beetle, Bruchidius siliquastri Delobel, 2007. *Wellcome Open Res*, *9*, 142. https://doi.org/10.12688/wellcomeopenres.21109.1

Barclay, M. V. L., Geiser, M., Vassiliades, D., Bayfield Farrell, W., Cristovao, J., Natural History Museum Genome Acquisition, L., Darwin Tree of Life Barcoding, c., Wellcome Sanger Institute Tree of Life Management, S., Laboratory, t., Wellcome Sanger Institute Scientific Operations: Sequencing, O., Wellcome Sanger Institute Tree of Life Core Informatics, t., Tree of Life Core Informatics, c., & Darwin Tree of Life, C. (2023). The genome sequence of a ground beetle, Pterostichus niger (Schaller, 1783). *Wellcome Open Res*, *8*, 544. https://doi.org/10.12688/wellcomeopenres.20418.1

Barclay, M. V. L., Geiser, M. F., Natural History Museum Genome Acquisition, L., Darwin Tree of Life Barcoding, c., Wellcome Sanger Institute Tree of Life Management, S., Laboratory, t., Wellcome Sanger Institute Scientific Operations: Sequencing, O., Wellcome Sanger Institute Tree of Life Core Informatics, t., Tree of Life Core Informatics, c., & Darwin Tree of Life, C. (2025). The genome sequence of a soldier beetle, Rhagonycha lutea (Muller, 1764). *Wellcome Open Res*, *10*, 24. https://doi.org/10.12688/wellcomeopenres.23467.1

Barclay, M. V. L., Namintraporn, T., Geiser, M. F., Natural History Museum Genome

Acquisition, L., Darwin Tree of Life Barcoding, c., Wellcome Sanger Institute Tree of Life Management, S., Laboratory, t., Wellcome Sanger Institute Scientific Operations: Sequencing, O., Wellcome Sanger Institute Tree of Life Core Informatics, t., Tree of Life Core Informatics, c., & Darwin Tree of Life, C. (2025). The genome sequence of 18-spot Ladybird, Myrrha octodecimguttata (Linnaeus, 1758). *Wellcome Open Res*, *10*, 7. https://doi.org/10.12688/wellcomeopenres.23455.1

Barclay, M. V. L., Natural History Museum Genome Acquisition, L., Darwin Tree of Life Barcoding, c., Wellcome Sanger Institute Tree of Life Management, S., Laboratory, t., Wellcome Sanger Institute Scientific Operations: Sequencing, O., Wellcome Sanger Institute Tree of Life Core Informatics, t., Tree of Life Core Informatics, c., & Darwin Tree of Life, C. (2024). The genome sequence of a soldier beetle, Cantharis flavilabris Fallen, 1807. *Wellcome Open Res*, *9*, 303. https://doi.org/10.12688/wellcomeopenres.22422.2

Barclay, M. V. L., Natural History Museum Genome Acquisition, L., Darwin Tree of Life Barcoding, c., Wellcome Sanger Institute Tree of Life, p., Wellcome Sanger Institute Scientific Operations, D. N. A. P. c., Tree of Life Core Informatics, c., & Darwin Tree of Life, C. (2023). The genome sequence of a ground beetle, Leistus spinibarbis (Fabricius, 1775). *Wellcome Open Res*, *8*, 412. https://doi.org/10.12688/wellcomeopenres.19997.1

Barclay, M. V. L., Natural History Museum Genome Acquisition, L., Darwin Tree of Life Barcoding, c., Wellcome Sanger Institute Tree of Life, p., Wellcome Sanger Institute Scientific Operations: Sequencing Operations, c., Tree of Life Core Informatics, c., & Darwin Tree of Life, C. (2023). The genome sequence of a weevil, Polydrusus cervinus (Linnaeus, 1758). *Wellcome Open Res*, *8*, 563. https://doi.org/10.12688/wellcomeopenres.20414.1

Barclay, M. V. L., Nikolaeva, S., Telnov, D., Natural History Museum Genome Acquisition, L., Darwin Tree of Life Barcoding, c., Wellcome Sanger Institute Tree of Life Management, S., Laboratory, t., Wellcome Sanger Institute Scientific Operations: Sequencing, O., Wellcome Sanger Institute Tree of Life Core Informatics, t., Tree of Life Core Informatics, c., & Darwin Tree of Life, C. (2025). The genome sequence of the false flower beetle, Anaspis frontalis (Linnaeus, 1758). *Wellcome Open Res*, *10*, 82. https://doi.org/10.12688/wellcomeopenres.23726.1

Barclay, M. V. L., Read, F., Namintraporn, T., Geiser, M. F., Natural History Museum Genome Acquisition, L., Darwin Tree of Life Barcoding, c., Wellcome Sanger Institute Tree of Life Management, S., Laboratory, t., Wellcome Sanger Institute Scientific Operations: Sequencing, O., Wellcome Sanger Institute Tree of Life Core Informatics, t., Tree of Life Core Informatics, c., & Darwin Tree of Life, C. (2025). The genome sequence of a beetle, Brachypterus glaber (Newman, 1834). *Wellcome Open Res*, *10*, 10. https://doi.org/10.12688/wellcomeopenres.23459.1

Barclay, M. V. L., Telnov, D., Natural History Museum Genome Acquisition, L., Darwin Tree of Life Barcoding, c., Wellcome Sanger Institute Tree of Life Management, S., Laboratory, t., Wellcome Sanger Institute Scientific Operations: Sequencing, O., Wellcome Sanger Institute Tree of Life Core Informatics, t., Tree of Life Core Informatics, c., & Darwin Tree of Life, C. (2025). The genome sequence of the acute-angled fungus beetle, Cryptophagus acutangulus Gyllenhal, 1827. *Wellcome Open Res*, *10*, 58. https://doi.org/10.12688/wellcomeopenres.23688.1

Barclay, M. V. L., Telnov, D., Natural History Museum Genome Acquisition, L., Darwin Tree of

Life Barcoding, c., Wellcome Sanger Institute Tree of Life Management, S., Laboratory, t., Wellcome Sanger Institute Scientific Operations: Sequencing, O., Wellcome Sanger Institute Tree of Life Core Informatics, t., Tree of Life Core Informatics, c., & Darwin Tree of Life, C. (2025). The genome sequence of the false flower beetle, Anaspis regimbarti Schilsky, 1895. *Wellcome Open Res*, *10*, 85. https://doi.org/10.12688/wellcomeopenres.23737.1

Barclay, M. V. L., Turner, C. R., Natural History Museum Genome Acquisition, L., Darwin Tree of Life Barcoding, c., Wellcome Sanger Institute Tree of Life Management, S., Laboratory, t., Wellcome Sanger Institute Scientific Operations: Sequencing, O., Wellcome Sanger Institute Tree of Life Core Informatics, t., Tree of Life Core Informatics, c., & Darwin Tree of Life, C. (2024). The genome sequence of a ground beetle, Harpalus rufipes (DeGeer, 1774). *Wellcome Open Res*, *9*, 483. https://doi.org/10.12688/wellcomeopenres.22875.1

Barclay, M. V. L., Turner, T., Natural History Museum Genome Acquisition, L., Darwin Tree of Life Barcoding, c., Wellcome Sanger Institute Tree of Life Management, S., Laboratory, t., Wellcome Sanger Institute Scientific Operations: Sequencing, O., Wellcome Sanger Institute Tree of Life Core Informatics, t., Tree of Life Core Informatics, c., & Darwin Tree of Life, C. (2025). The genome sequence of a seed weevil, Aspidapion aeneum (Fabricius, 1775). *Wellcome Open Res*, *10*, 41. https://doi.org/10.12688/wellcomeopenres.23613.1

Barclay, M. V. L., Turner, T., Natural History Museum Genome Acquisition Lab, Darwin Tree of Life Barcoding collective, Wellcome Sanger Institute Tree of Life Management, S. a. L., team, Wellcome Sanger Institute Scientific Operations: Sequencing Operations, Wellcome Sanger Institute Tree of Life Core Informatics team, & Tree of Life Core Informatics collective, D. T. o. L. C. (2025). The genome sequence of a Ground Beetle, Harpalus Rubripes (Duftschmid, 1812). *Wellcome Open Res*. https://doi.org/10.12688/wellcomeopenres.23433.1

Barclay, M. V. L., Turner, T., Telfer, M. G., Geiser, M. F., Natural History Museum Genome Acquisition, L., University of, O., Wytham Woods Genome Acquisition, L., Darwin Tree of Life Barcoding, c., Wellcome Sanger Institute Tree of Life Management, S., Laboratory, t., Wellcome Sanger Institute Scientific Operations: Sequencing, O., Wellcome Sanger Institute Tree of Life Core Informatics, t., Tree of Life Core Informatics, c., & Darwin Tree of Life, C. (2025). The genome sequence of a bark-dwelling beetle, Silvanus unidentatus (Olivier, 1790). *Wellcome Open Res*, *10*, 136. https://doi.org/10.12688/wellcomeopenres.23771.1

Barclay, M. V. L., Vassiliades, D., Bayfield Farrell, W., Cristovao, J., Matsumoto, K., Geiser, M., Telfer, M. G., Natural History Museum Genome Acquisition, L., University of, O., Wytham Woods Genome Acquisition, L., Darwin Tree of Life Barcoding, c., Wellcome Sanger Institute Tree of Life Management, S., Laboratory, t., Wellcome Sanger Institute Scientific Operations: Sequencing, O., Wellcome Sanger Institute Tree of Life Core Informatics, t., Tree of Life Core Informatics, c., & Darwin Tree of Life, C. (2024). The genome sequence of the oak pinhole borer, Platypus cylindrus (Fabricius, 1792). *Wellcome Open Res*, *9*, 305. https://doi.org/10.12688/wellcomeopenres.22425.2

Barclay, M. V. L., Vassiliades, D., Bayfield-Farrell, W., Cristovao, J., Matsumoto, K., Geiser, M., Natural History Museum Genome Acquisition, L., Darwin Tree of Life Barcoding, c., Wellcome Sanger Institute Tree of Life Management, S., Laboratory, t., Wellcome

Sanger Institute Scientific Operations: Sequencing, O., Wellcome Sanger Institute Tree of Life Core Informatics, t., Tree of Life Core Informatics, c., & Darwin Tree of Life, C. (2024). The genome sequence of the willow leaf beetle, Lochmaea capreae Linnaeus, 1758. *Wellcome Open Res*, *9*, 304. https://doi.org/10.12688/wellcomeopenres.22424.2

Bolanakis, G., Karakasi, D., Trichas, A., Bohne, A., Monteiro, R., Fernandez, R., Escudero, N., Bitzilekis, E., Stratakis, M., Lymberakis, P., Poulakakis, N., Genoscope Sequencing, T., Moussy, A., Cruaud, C., Labadie, K., Demirdjian, L., Duprat, S., Teodori, E., Wincker, P., . . . Bortoluzzi, C. (2025). ERGA-BGE genome of Dendarus foraminosus: an IUCN Least Concern darkling beetle endemic to Crete (Greece). *Open Res Eur*, *5*, 173. https://doi.org/10.12688/openreseurope.20489.3

Booth, R., Crowley, L. M., Natural History Museum Genome Acquisition, L., University of, O., Wytham Woods Genome Acquisition, L., Darwin Tree of Life Barcoding, C., Wellcome Sanger Institute Tree of Life Management, S., Laboratory, t., Wellcome Sanger Institute Scientific Operations: Sequencing, O., Wellcome Sanger Institute Tree of Life Core Informatics, t., Tree of Life Core Informatics, c., & Darwin Tree of Life, C. (2025). The genome sequence of a Chequered Beetle, Tillus elongatus (Linnaeus, 1758) (Coleoptera: Cleridae). *Wellcome Open Res*, *10*, 575. https://doi.org/10.12688/wellcomeopenres.24678.1

Booth, R., Darwin Tree of Life Barcoding collective, Wellcome Sanger Institute Tree of Life Management, S. a. L., team, Wellcome Sanger Institute Scientific Operations: Sequencing Operations, Wellcome Sanger Institute Tree of Life Core Informatics team, & Tree of Life Core Informatics collective, D. T. o. L. C. (2024). The genome sequence of a darkling beetle, Diaperis boleti (Linnaeus, 1758). *Wellcome Open Res*. https://doi.org/10.12688/wellcomeopenres.23071.1

Booth, R., Natural History Museum Genome Acquisition, L., Darwin Tree of Life Barcoding, c., Wellcome Sanger Institute Tree of Life Management, S., Laboratory, t., Wellcome Sanger Institute Scientific Operations: Sequencing, O., Wellcome Sanger Institute Tree of Life Core Informatics, t., Tree of Life Core Informatics, c., & Darwin Tree of Life, C. (2024). The genome sequence of a fungus weevil, Pseudeuparius sepicola (Fabricius, 1792). *Wellcome Open Res*, *9*, 615. https://doi.org/10.12688/wellcomeopenres.23209.1

Booth, R., Natural History Museum Genome Acquisition, L., Darwin Tree of Life Barcoding, c., Wellcome Sanger Institute Tree of Life Management, S., Laboratory, t., Wellcome Sanger Institute Scientific Operations: Sequencing, O., Wellcome Sanger Institute Tree of Life Core Informatics, t., Tree of Life Core Informatics, c., & Darwin Tree of Life, C. (2024). The genome sequence of a leaf beetle, Galeruca laticollis Sahlberg, C.R., 1838. *Wellcome Open Res*, *9*, 594. https://doi.org/10.12688/wellcomeopenres.23195.1

Booth, R., Natural History Museum Genome Acquisition, L., Darwin Tree of Life Barcoding, c., Wellcome Sanger Institute Tree of Life Management, S., Laboratory, t., Wellcome Sanger Institute Scientific Operations: Sequencing, O., Wellcome Sanger Institute Tree of Life Core Informatics, t., Tree of Life Core Informatics, c., & Darwin Tree of Life, C. (2025). The genome sequence of a leaf beetle, Galerucella nymphaeae (Linnaeus, 1758). *Wellcome Open Res*, *10*, 57. https://doi.org/10.12688/wellcomeopenres.23676.1

Boyes, D., Crowley, L. M., Holland, P. W. H., University of, O., Wytham Woods Genome Acquisition, L., Darwin Tree of Life Barcoding, c., Wellcome Sanger Institute Tree of Life Management, S., Laboratory, t., Wellcome Sanger Institute Scientific Operations: Sequencing, O., Wellcome Sanger Institute Tree of Life Core Informatics, t., Tree of Life

Core Informatics, c., & Darwin Tree of Life, C. (2024). The genome sequence of the Summer Chafer, Amphimallon solstitiale (Linnaeus, 1758). *Wellcome Open Res*, *9*, 138. https://doi.org/10.12688/wellcomeopenres.21100.1

Bracewell, R. R., Stillman, J. H., Dahlhoff, E. P., Smeds, E., Chatla, K., Bachtrog, D., Williams, C., & Rank, N. E. (2023). A chromosome-scale genome assembly and evaluation of mtDNA variation in the willow leaf beetle Chrysomela aeneicollis. *G3 (Bethesda)*, *13*(7). https://doi.org/10.1093/g3journal/jkad106

Chen, M., Mei, Y., Chen, X., Chen, X., Xiao, D., He, K., Li, Q., Wu, M., Wang, S., Zhang, F., & Li, F. (2021). A chromosome-level assembly of the harlequin ladybird Harmonia axyridis as a genomic resource to study beetle and invasion biology. *Mol Ecol Resour*, *21*(4), 1318-1332. https://doi.org/10.1111/1755-0998.13342

Cohen, Z. P., Perkin, L. C., Sim, S. B., Stahlke, A. R., Geib, S. M., Childers, A. K., Smith, T. P. L., & Suh, C. (2023). Insight into weevil biology from a reference quality genome of the boll weevil, Anthonomus grandis grandis Boheman (Coleoptera: Curculionidae). *G3 (Bethesda)*, *13*(2). https://doi.org/10.1093/g3journal/jkac309

Copeland, M., Landa, S., Owoyemi, A. O., Jonika, M. M., Alfieri, J. M., Johnston, J. S., Sylvester, T. P., Kyre, B. R., Hoover, Z., Hjelmen, C. E., Rieske, L. K., Blackmon, H., & Casola, C. (2024). Genome assembly of the southern pine beetle (Dendroctonus frontalis Zimmerman) reveals the origins of gene content reduction in Dendroctonus. *R Soc Open Sci*, *11*(12), 240755. https://doi.org/10.1098/rsos.240755

Crellin, S., Natural History Museum Genome Acquisition, L., Darwin Tree of Life Barcoding, C., Wellcome Sanger Institute Tree of Life Management, S., Laboratory, t., Wellcome Sanger Institute Scientific Operations: Sequencing, O., Wellcome Sanger Institute Tree of Life Core Informatics, t., Tree of Life Core Informatics, c., & Darwin Tree of Life, C. (2025). The genome sequence of a longhorn beetle, Stictoleptura scutellata (Fabricius, 1781) (Coleoptera: Cerambycidae). *Wellcome Open Res*, *10*, 615. https://doi.org/10.12688/wellcomeopenres.25051.1

Crowley, L., McCulloch, J., University of Oxford and Wytham Woods Genome Acquisition Lab, Darwin Tree of Life Barcoding collective, Wellcome Sanger Institute Tree of Life programme, Wellcome Sanger Institute Scientific Operations: Sequencing Operations, & Tree of Life Core Informatics collective, D. T. o. L. C. (2023). The genome sequence of a soldier beetle, Cantharis nigra (DeGeer, 1774). *Wellcome Open Res*. https://doi.org/10.12688/wellcomeopenres.20382.1

Crowley, L., University of, O., Wytham Woods Genome Acquisition, L., Darwin Tree of Life Barcoding, c., Wellcome Sanger Institute Tree of Life, p., Wellcome Sanger Institute Scientific Operations, D. N. A. P. c., Tree of Life Core Informatics, c., & Darwin Tree of Life, C. (2021). The genome sequence of the seven-spotted ladybird, Coccinella septempunctata Linnaeus, 1758. *Wellcome Open Res*, *6*, 319. https://doi.org/10.12688/wellcomeopenres.17346.1

Crowley, L., University of, O., Wytham Woods Genome Acquisition, L., Darwin Tree of Life Barcoding, c., Wellcome Sanger Institute Tree of Life, p., Wellcome Sanger Institute Scientific Operations, D. N. A. P. c., Tree of Life Core Informatics, c., Garner, B., & Darwin Tree of Life, C. (2023). The genome sequence of a ground beetle, Nebria brevicollis (Fabricius, 1792). *Wellcome Open Res*, *8*, 20. https://doi.org/10.12688/wellcomeopenres.18749.1

Crowley, L., University of Oxford and Wytham Woods Genome Acquisition Lab, Darwin Tree of Life Barcoding collective, Wellcome Sanger Institute Tree of Life Management, S. a. L., team, Wellcome Sanger Institute Scientific Operations: Sequencing Operations, Wellcome Sanger Institute Tree of Life Core Informatics team, & Tree of Life Core Informatics collective, D. T. o. L. C. (2025). The genome sequence of the brown spruce longhorn beetle Tetropium fuscum (Fabricius, 1787). *Wellcome Open Res*. https://doi.org/10.12688/wellcomeopenres.24296.1

Crowley, L., University of Oxford and Wytham Woods Genome Acquisition Lab, Darwin Tree of Life Barcoding collective, Wellcome Sanger Institute Tree of Life programme, Wellcome Sanger Institute Scientific Operations: DNA Pipelines collective, & Tree of Life Core Informatics collective, D. T. o. L. C. (2021). The genome sequence of the hazel leaf-roller, Apoderus coryli (Linnaeus, 1758). *Wellcome Open Res*. https://doi.org/10.12688/wellcomeopenres.17380.1

Crowley, L., Whiffin, A., University of Oxford and Wytham Woods Genome Acquisition Lab, Darwin Tree of Life Barcoding collective, Wellcome Sanger Institute Tree of Life programme, Wellcome Sanger Institute Scientific Operations: DNA Pipelines collective, Wellcome Sanger Institute Tree of Life Core Informatics Team, & Tree of Life Core Informatics collective, D. T. o. L. C. (2023). The genome sequence of the Common Snail-hunter beetle, Phosphuga atrata (Linnaeus, 1758). *Wellcome Open Res*. https://doi.org/10.12688/wellcomeopenres.20340.1

Crowley, L. M., Badham, X. R., University of, O., Wytham Woods Genome Acquisition, L., Darwin Tree of Life Barcoding, c., Wellcome Sanger Institute Tree of Life Management, S., Laboratory, t., Wellcome Sanger Institute Scientific Operations: Sequencing, O., Wellcome Sanger Institute Tree of Life Core Informatics, t., Tree of Life Core Informatics, c., & Darwin Tree of Life, C. (2025). The genome sequence of a fungus weevil, Platystomos albinus (C.Linnaeus, 1758). *Wellcome Open Res*, *10*, 295. https://doi.org/10.12688/wellcomeopenres.24290.1

Crowley, L. M., Barclay, M., Roy, H. E., Brown, P. M. J., University of, O., Wytham Woods Genome Acquisition, L., Natural History Museum Genome Acquisition, L., Darwin Tree of Life Barcoding, c., Wellcome Sanger Institute Tree of Life, p., Wellcome Sanger Institute Scientific Operations, D. N. A. P. c., Tree of Life Core Informatics, c., & Darwin Tree of Life, C. (2023). The genome sequence of the orange ladybird, Halyzia sedecimguttata (Linnaeus, 1758). *Wellcome Open Res*, *8*, 186. https://doi.org/10.12688/wellcomeopenres.19369.1

Crowley, L. M., Booth, R., Geiser, M. F., University of, O., Wytham Woods Acquisition, L., Darwin Tree of Life Barcoding, C., Wellcome Sanger Institute Tree of Life Management, S., Laboratory, t., Wellcome Sanger Institute Scientific Operations: Sequencing, O., Wellcome Sanger Institute Tree of Life Core Informatics, t., Tree of Life Core Informatics, c., & Darwin Tree of Life, C. (2025). The genome sequence of a ground beetle, Dromius quadrimaculatus (Linnaeus, 1758) (Coleoptera: Carabidae). *Wellcome Open Res*, *10*, 555. https://doi.org/10.12688/wellcomeopenres.24898.1

Crowley, L. M., Broad, G. R., Fletcher, C., Januszczak, I., Barnes, I., Whiffin, A. L., Natural History Museum Genome Acquisition, L., Darwin Tree of Life Barcoding, c., Wellcome Sanger Institute Tree of Life Management, S., Laboratory, t., Wellcome Sanger Institute Scientific Operations: Sequencing, O., Wellcome Sanger Institute Tree of Life Core Informatics, t., Tree of Life Core Informatics, c., & Darwin Tree of Life, C. (2024). The genome sequence of the Banded Burying beetle, Nicrophorus investigator Zetterstedt,

1824. *Wellcome Open Res*, *9*, 343. https://doi.org/10.12688/wellcomeopenres.21496.1
Crowley, L. M., McCulloch, J., University of, O., Wytham Woods Genome Acquisition, L., Darwin Tree of Life Barcoding, C., Wellcome Sanger Institute Tree of Life Management, S., Laboratory, t., Wellcome Sanger Institute Scientific Operations: Sequencing, O., Wellcome Sanger Institute Tree of Life Core Informatics, t., Tree of Life Core Informatics, c., & Darwin Tree of Life, C. (2026). The genome sequence of the soldier beetle, Malthodes minimus (Linnaeus, 1758) (Coleoptera: Cantharidae). *Wellcome Open Res*, *11*, 16. https://doi.org/10.12688/wellcomeopenres.25397.1
Crowley, L. M., Phillips, D., University of, O., Wytham Woods Genome Acquisition, L., Darwin Tree of Life Barcoding, c., Wellcome Sanger Institute Tree of Life Management, S., Laboratory, t., Wellcome Sanger Institute Scientific Operations: Sequencing, O., Wellcome Sanger Institute Tree of Life Core Informatics, t., Tree of Life Core Informatics, c., & Darwin Tree of Life, C. (2024). The genome sequence of the lesser stag beetle, Dorcus parallelipipedus (Linnaeus, 1758). *Wellcome Open Res*, *9*, 202. https://doi.org/10.12688/wellcomeopenres.21262.1
Crowley, L. M., Roy, H. E., Brown, P. M. J., University of, O., Wytham Woods Genome Acquisition, L., Darwin Tree of Life Barcoding, c., Wellcome Sanger Institute Tree of Life Management, S., Laboratory, t., Wellcome Sanger Institute Scientific Operations: Sequencing, O., Wellcome Sanger Institute Tree of Life Core Informatics, t., Tree of Life Core Informatics, c., & Darwin Tree of Life, C. (2024). The genome sequence of the ten-spot ladybird, Adalia decempunctata (Linnaeus, 1758). *Wellcome Open Res*, *9*, 106. https://doi.org/10.12688/wellcomeopenres.21008.1
Crowley, L. M., Sudworth, J., University of, O., Wytham Woods Genome Acquisition, L., Darwin Tree of Life Barcoding, c., Wellcome Sanger Institute Tree of Life, p., Wellcome Sanger Institute Scientific Operations, D. N. A. P. c., Tree of Life Core Informatics, c., & Darwin Tree of Life, C. (2023). The genome sequence of a ground beetle, Ophonus ardosiacus (Lutshnik, 1922). *Wellcome Open Res*, *8*, 353. https://doi.org/10.12688/wellcomeopenres.19849.1
Crowley, L. M., Telfer, M., Barclay, M. V. L., Badham, X. R., University of, O., Wytham Woods Genome Acquisition, L., Natural History Museum Genome Acquisition, L., Darwin Tree of Life Barcoding, c., Wellcome Sanger Institute Tree of Life Management, S., Laboratory, t., Wellcome Sanger Institute Scientific Operations: Sequencing, O., Wellcome Sanger Institute Tree of Life Core Informatics, t., Tree of Life Core Informatics, c., & Darwin Tree of Life, C. (2024). The genome sequence of a ground beetle, Clivina fossor (Linnaeus, 1758). *Wellcome Open Res*, *9*, 438. https://doi.org/10.12688/wellcomeopenres.22898.1
Crowley, L. M., Telfer, M., Barclay, M. V. L., Phillips, D., University of, O., Wytham Woods Genome Acquisition, L., Natural History Museum Genome Acquisition, L., Darwin Tree of Life Barcoding, c., Wellcome Sanger Institute Tree of Life Management, S., Laboratory, t., Wellcome Sanger Institute Scientific Operations: Sequencing, O., Wellcome Sanger Institute Tree of Life Core Informatics, t., Tree of Life Core Informatics, c., & Darwin Tree of Life, C. (2024). The genome sequence of the Dogs-Mercury Flea Beetle, Hermaeophaga mercurialis (Fabricius, 1792). *Wellcome Open Res*, *9*, 503. https://doi.org/10.12688/wellcomeopenres.22896.1
Crowley, L. M., Telfer, M., Escalona, H. E., University of Oxford and Wytham Woods Genome Acquisition Lab, Darwin Tree of Life Barcoding collective, Wellcome Sanger Institute

Tree of Life Management, S. a. L., team, Wellcome Sanger Institute Scientific Operations: Sequencing Operations, Wellcome Sanger Institute Tree of Life Core Informatics team, & Tree of Life Core Informatics collective, D. T. o. L. C. (2024). The genome sequence of a ground beetle, Agonum fuliginosum (Panzer, 1809). *Wellcome Open Res*. https://doi.org/10.12688/wellcomeopenres.20912.1

Crowley, L. M., Telfer, M., Geiser, M., University of, O., Wytham Woods Genome Acquisition, L., Mulley, J. F., Natural History Museum Genome Acquisition, L., Darwin Tree of Life Barcoding, c., Wellcome Sanger Institute Tree of Life, p., Wellcome Sanger Institute Scientific Operations, D. N. A. P. c., Tree of Life Core Informatics, c., & Darwin Tree of Life, C. (2023). The genome sequence of Philonthus cognatus (Stephens, 1832) (Coleoptera, Staphylinidae), a rove beetle. *Wellcome Open Res*, *8*, 169. https://doi.org/10.12688/wellcomeopenres.19336.1

Crowley, L. M., Telfer, M., McCulloch, J., University of Oxford and Wytham Woods Genome Acquisition Lab, Darwin Tree of Life Barcoding collective, Wellcome Sanger Institute Tree of Life Management, S. a. L., team, Wellcome Sanger Institute Scientific Operations: Sequencing Operations, Wellcome Sanger Institute Tree of Life Core Informatics team, & Tree of Life Core Informatics collective, D. T. o. L. C. (2025). The genome sequence of a rove beetle, Stenus bimaculatus (Gyllenhal, 1810). *Wellcome Open Res*. https://doi.org/10.12688/wellcomeopenres.24262.1

Crowley, L. M., Telfer, M., University of Oxford and Wytham Woods Genome Acquisition Lab, Darwin Tree of Life Barcoding collective, Wellcome Sanger Institute Tree of Life Management, S. a. L., team, Wellcome Sanger Institute Scientific Operations: Sequencing Operations, Wellcome Sanger Institute Tree of Life Core Informatics team, & Tree of Life Core Informatics collective, D. T. o. L. C. (2025). The genome sequence of the flat bark beetle, Pediacus dermestoides (Fabricius, 1792). *Wellcome Open Res*. https://doi.org/10.12688/wellcomeopenres.24345.1

Crowley, L. M., Telfer, M. G., Escalona, H. E., University of, O., Wytham Woods Genome Acquisition, L., Darwin Tree of Life Barcoding, c., Wellcome Sanger Institute Tree of Life Management, S., Laboratory, t., Wellcome Sanger Institute Scientific Operations: Sequencing, O., Wellcome Sanger Institute Tree of Life Core Informatics, t., Tree of Life Core Informatics, c., & Darwin Tree of Life, C. (2024). The genome sequence of the hawthorn leaf beetle, Lochmaea crataegi (Forster, 1771). *Wellcome Open Res*, *9*, 80. https://doi.org/10.12688/wellcomeopenres.20911.1

Crowley, L. M., University of, O., Wytham Woods Genome Acquisition, L., Darwin Tree of Life Barcoding, C., Wellcome Sanger Institute Tree of Life Management, S., Laboratory, t., Wellcome Sanger Institute Scientific Operations: Sequencing, O., Wellcome Sanger Institute Tree of Life Core Informatics, t., Tree of Life Core Informatics, c., & Darwin Tree of Life, C. (2026). The genome sequence of the cryptorhynchine weevil, Kyklioacalles roboris (Stuben, 2003) (Coleoptera: Curculionidae). *Wellcome Open Res*, *11*, 11. https://doi.org/10.12688/wellcomeopenres.25416.1

Crowley, L. M., University of, O., Wytham Woods Genome Acquisition, L., Darwin Tree of Life Barcoding, c., Wellcome Sanger Institute Tree of Life, p., Wellcome Sanger Institute Scientific Operations, D. N. A. P. c., Tree of Life Core Informatics, c., Chua, P., Kusy, D., & Darwin Tree of Life, C. (2023). The genome sequence of a soldier beetle, Podabrus alpinus (Paykull, 1798). *Wellcome Open Res*, *8*, 56. https://doi.org/10.12688/wellcomeopenres.18890.1

Crowley, L. M., University of, O., Wytham Woods Genome Acquisition, L., Darwin Tree of Life Barcoding, c., Wellcome Sanger Institute Tree of Life, p., Wellcome Sanger Institute Scientific Operations, D. N. A. P. c., Tree of Life Core Informatics, c., & Darwin Tree of Life, C. (2021). The genome sequence of the black clock beetle, Pterostichus madidus (Fabricius, 1775). *Wellcome Open Res*, *6*, 301. https://doi.org/10.12688/wellcomeopenres.17347.1

Crowley, L. M., University of, O., Wytham Woods Genome Acquisition, L., Darwin Tree of Life Barcoding, c., Wellcome Sanger Institute Tree of Life, p., Wellcome Sanger Institute Scientific Operations, D. N. A. P. c., Tree of Life Core Informatics, c., & Darwin Tree of Life, C. (2021). The genome sequence of the common malachite beetle, Malachius bipustulatus (Linnaeus, 1758). *Wellcome Open Res*, *6*, 322. https://doi.org/10.12688/wellcomeopenres.17381.1

Crowley, L. M., University of, O., Wytham Woods Genome Acquisition, L., Darwin Tree of Life Barcoding, c., Wellcome Sanger Institute Tree of Life, p., Wellcome Sanger Institute Scientific Operations, D. N. A. P. c., Tree of Life Core Informatics, c., & Darwin Tree of Life, C. (2021). The genome sequence of the devil's coach horse beetle, Ocypus olens (Muller, 1764). *Wellcome Open Res*, *6*, 293. https://doi.org/10.12688/wellcomeopenres.17342.2

Crowley, L. M., University of, O., Wytham Woods Genome Acquisition, L., Natural History Museum Genome Acquisition, L., Darwin Tree of Life Barcoding, c., Wellcome Sanger Institute Tree of Life, p., Wellcome Sanger Institute Scientific Operations, D. N. A. P. c., Tree of Life Core Informatics, c., & Darwin Tree of Life, C. (2021). The genome sequence of the common red soldier beetle, Rhagonycha fulva (Scopoli, 1763). *Wellcome Open Res*, *6*, 243. https://doi.org/10.12688/wellcomeopenres.17198.1

Delic, T., Wellcome Sanger Institute Tree of Life Management, S., Laboratory, t., Wellcome Sanger Institute Scientific Operations: Sequencing, O., Tree of Life Core Informatics, c., & Wellcome Sanger Institute Tree of Life Core Informatics, t. (2025). The genome sequence of a cave beetle, Leptodirus hochenwartii F.J.Schmidt, 1832. *Wellcome Open Res*, *10*, 159. https://doi.org/10.12688/wellcomeopenres.23959.1

Findlay, J. D. S., Foster, G., Natural History Museum Genome Acquisition, L., Darwin Tree of Life Barcoding, c., Wellcome Sanger Institute Tree of Life, p., Wellcome Sanger Institute Scientific Operations, D. N. A. P. c., Tree of Life Core Informatics, c., & Darwin Tree of Life, C. (2023). The genome sequence of a riffle beetle, Elmis aenea (Muller, 1806). *Wellcome Open Res*, *8*, 322. https://doi.org/10.12688/wellcomeopenres.19778.1

Fu, N., Li, J., Ren, L., Li, X., Wang, M., Li, F., Zong, S., & Luo, Y. (2022). Chromosome-level genome assembly of Monochamus saltuarius reveals its adaptation and interaction mechanism with pine wood nematode. *Int J Biol Macromol*, *222*(Pt A), 325-336. https://doi.org/10.1016/j.ijbiomac.2022.09.108

Gao, Y. F., Yang, F. Y., Song, W., Cao, L. J., Chen, J. C., Shen, X. J., Qu, L. J., Zong, S. X., & Wei, S. J. (2024). Chromosome-level genome assembly of the Japanese sawyer beetle Monochamus alternatus. *Sci Data*, *11*(1), 199. https://doi.org/10.1038/s41597-024-03048-y

Garland, S., Natural History Museum Genome Acquisition, L., Darwin Tree of Life Barcoding, c., Wellcome Sanger Institute Tree of Life Management, S., Laboratory, t., Wellcome Sanger Institute Scientific Operations: Sequencing, O., Wellcome Sanger Institute Tree of Life Core Informatics, t., Tree of Life Core Informatics, c., & Darwin Tree of Life, C.

(2024). The genome sequence of a jewel beetle, Agrilus biguttatus (Fabricius, 1776). *Wellcome Open Res*, *9*, 413. https://doi.org/10.12688/wellcomeopenres.22762.1

Geiser, M. F., Read, F., Barclay, M. V. L., Natural History Museum Genome Acquisition, L., Darwin Tree of Life Barcoding, c., Wellcome Sanger Institute Tree of Life Management, S., Laboratory, t., Wellcome Sanger Institute Scientific Operations: Sequencing, O., Wellcome Sanger Institute Tree of Life Core Informatics, t., Tree of Life Core Informatics, c., & Darwin Tree of Life, C. (2025). The genome sequence of the Asparagus Beetle, Crioceris asparagi (Linnaeus, 1758). *Wellcome Open Res*, *10*, 18. https://doi.org/10.12688/wellcomeopenres.23460.1

Geiser, M. F., Read, F., Barclay, M. V. L., Natural History Museum Genome Acquisition Lab, Darwin Tree of Life Barcoding collective, Wellcome Sanger Institute Tree of Life Management, S. a. L., team, Wellcome Sanger Institute Scientific Operations: Sequencing Operations, Wellcome Sanger Institute Tree of Life Core Informatics team, & Tree of Life Core Informatics collective, D. T. o. L. C. (2025). The genome sequence of the Rosemary Beetle, Chrysolina americana (Linnaeus, 1758). *Wellcome Open Res*. https://doi.org/10.12688/wellcomeopenres.23458.1

Geiser, M. F., Sims, I., Natural History Museum Genome Acquisition, L., Darwin Tree of Life Barcoding, c., Wellcome Sanger Institute Tree of Life Management, S., Laboratory, t., Wellcome Sanger Institute Scientific Operations: Sequencing, O., Wellcome Sanger Institute Tree of Life Core Informatics, t., Tree of Life Core Informatics, c., & Darwin Tree of Life, C. (2025). The genome sequence of a flea beetle, Neocrepidodera transversa (Marsham, 1802). *Wellcome Open Res*, *10*, 62. https://doi.org/10.12688/wellcomeopenres.23697.1

Grayson, A., Geiser, M., Natural History Museum Genome Acquisition Lab, Darwin Tree of Life Barcoding collective, Wellcome Sanger Institute Tree of Life programme, Operations, W. S. I. S. O. S., collective, & Tree of Life Core Informatics collective, D. T. o. L. C. (2023). The genome sequence of the Rose Chafer, Cetonia aurata (Linnaeus, 1758). *Wellcome Open Res*. https://doi.org/10.12688/wellcomeopenres.20412.1

Hislop, L. W. S. I. T. o. L. M., Samples and Laboratory, team, Wellcome Sanger Institute Scientific Operations: Sequencing Operations, Wellcome Sanger Institute Tree of Life Core Informatics team, & Tree of Life Core Informatics collective, D. T. o. L. C. (2024). The genome sequence of an elaterid beetle, Ctenicera cuprea (Fabricius, 1775). *Wellcome Open Res*. https://doi.org/10.12688/wellcomeopenres.22885.1

Jefferys, E. E., Holland, P. W. H., Thomas, P., Hugman, M., University of, O., Wytham Woods Genome Acquisition, L., Darwin Tree of Life Barcoding, c., Wellcome Sanger Institute Tree of Life Management, S., Laboratory, t., Wellcome Sanger Institute Scientific Operations: Sequencing, O., Wellcome Sanger Institute Tree of Life Core Informatics, t., Tree of Life Core Informatics, c., & Darwin Tree of Life, C. (2024). The genome sequence of the Deathwatch beetle, Xestobium rufovillosum (De Geer, 1774). *Wellcome Open Res*, *9*, 619. https://doi.org/10.12688/wellcomeopenres.23210.1

Jin, J., Zhan, Z., Ye, M., & Jing, S. (2024). A chromosomal-level genome assembly of Serrognathus titanus Boisduval, 1835 (Coleoptera: Lucanidae). *Sci Data*, *11*(1), 888. https://doi.org/10.1038/s41597-024-03727-w

Levey, B., Natural History Museum Genome Acquisition, L., Darwin Tree of Life Barcoding, c., Wellcome Sanger Institute Tree of Life Management, S., Laboratory, t., Wellcome Sanger Institute Scientific Operations: Sequencing, O., Wellcome Sanger Institute Tree of Life Core Informatics, t., Tree of Life Core Informatics, c., & Darwin Tree of Life, C. (2024). The genome sequence of a leaf beetle, Chrysolina haemoptera (Linnaeus, 1758). *Wellcome Open Res*, *9*, 427. https://doi.org/10.12688/wellcomeopenres.22887.1

Li, X., Mao, C., He, J., Bin, X., Liu, G., Dong, Z., Zhao, R., Wan, X., & Li, X. (2024). The first chromosome-level genome of the stag beetle Dorcus hopei Saunders, 1854 (Coleoptera: Lucanidae). *Sci Data*, *11*(1), 396. https://doi.org/10.1038/s41597-024-03251-x

Liam M. Crowley, R. P., University of Oxford and Wytham Woods Genome Acquisition Lab, Darwin Tree of Life Barcoding collective, Wellcome Sanger Institute Tree of Life Management, S. a. L., team, Wellcome Sanger Institute Scientific Operations: Sequencing Operations, Wellcome Sanger Institute Tree of Life Core Informatics team, & Tree of Life Core Informatics collective, D. T. o. L. C. (2024). The genome sequence of a longhorn beetle, Stenurella melanura (Linnaeus, 1758). *Wellcome Open Res*. https://doi.org/10.12688/wellcomeopenres.22580.1

Liam M. Crowley, U. o. O. a. W. W. A. L., Darwin Tree of Life Barcoding Collective, Wellcome Sanger Institute Tree of Life Management, S. a. L., team, Wellcome Sanger Institute Scientific Operations: Sequencing Operations, Wellcome Sanger Institute Tree of Life Core Informatics team, & Tree of Life Core Informatics collective, D. T. o. L. C. (2025). The genome assembly of a soft-winged flower beetle, Anthocomus fasciatus (Linnaeus, 1758) (Coleoptera: Melyridae). *Wellcome Open Res*. https://doi.org/10.12688/wellcomeopenres.24976.1

Lyszkowski, R., Telnov, D., Barclay, M. V. L., Natural History Museum Genome Acquisition, L., Darwin Tree of Life Barcoding, c., Wellcome Sanger Institute Tree of Life Management, S., Laboratory, t., Wellcome Sanger Institute Scientific Operations: Sequencing, O., Wellcome Sanger Institute Tree of Life Core Informatics, t., Tree of Life Core Informatics, c., & Darwin Tree of Life, C. (2024). The genome sequence of the Scarce Cardinal Beetle, Schizotus pectinicornis (Linnaeus, 1758). *Wellcome Open Res*, *9*, 501. https://doi.org/10.12688/wellcomeopenres.22888.1

Mann, D. J., Crowley, L. M., Fletcher, C., Angus, R., Badham, X. R., University of, O., Wytham Woods Genome Acquisition, L., Natural History Museum Genome Acquisition, L., Darwin Tree of Life Barcoding, C., Wellcome Sanger Institute Tree of Life Management, S., Laboratory, t., Wellcome Sanger Institute Scientific Operations: Sequencing, O., Wellcome Sanger Institute Tree of Life Core Informatics, t., Tree of Life Core Informatics, c., & Darwin Tree of Life, C. (2025). The genome sequence of the common dung beetle, Aphodius fimetarius (Linnaeus, 1758) (Coleoptera: Scarabaeidae). *Wellcome Open Res*, *10*, 519. https://doi.org/10.12688/wellcomeopenres.24863.1

Mann, D. J., Crowley, L. M., Sivell, O., Booth, R., University of Oxford and Wytham Woods Genome Acquisition Lab, Natural History Museum Genome Acquisition Lab, Darwin Tree of Life Barcoding collective, Wellcome Sanger Institute Tree of Life Management, S. a. L., team, Wellcome Sanger Institute Scientific Operations: Sequencing Operations, Wellcome Sanger Institute Tree of Life Core Informatics team, & Tree of Life Core Informatics collective, D. T. o. L. C. (2025). The genome sequence of a soldier beetle, Cantharis lateralis (Linnaeus, 1758). *Wellcome Open Res*. https://doi.org/10.12688/wellcomeopenres.24002.1

Mann, D. J., Crowley, L. M., University of, O., Wytham Woods Genome Acquisition, L., Darwin Tree of Life Barcoding, c., Wellcome Sanger Institute Tree of Life Management, S., Laboratory, t., Wellcome Sanger Institute Scientific Operations: Sequencing, O.,

Wellcome Sanger Institute Tree of Life Core Informatics, t., Tree of Life Core Informatics, c., & Darwin Tree of Life, C. (2025). The genome sequence of a dung beetle, Aphodius ( Calamosternus) granarius Linnaeus, 1767. *Wellcome Open Res*, *10*, 369. https://doi.org/10.12688/wellcomeopenres.24306.2

Mann, D. J., Crowley, L. M., University of, O., Wytham Woods Genome Acquisition, L., Darwin Tree of Life Barcoding, c., Wellcome Sanger Institute Tree of Life Management, S., Laboratory, t., Wellcome Sanger Institute Scientific Operations: Sequencing, O., Wellcome Sanger Institute Tree of Life Core Informatics, t., Tree of Life Core Informatics, c., & Darwin Tree of Life, C. (2025). The genome sequence of a rove beetle, Philonthus spinipes Sharp, 1874. *Wellcome Open Res*, *10*, 217. https://doi.org/10.12688/wellcomeopenres.23996.1

Maxwell V. L. Barclay, M. F. G., Keita Matsumoto,, Danaë Vassiliades, J. C., Will Bayfield Farrell,, Natural History Museum Genome Acquisition Lab, Darwin Tree of Life Barcoding Collective, Wellcome Sanger Institute Tree of Life Management, S. a. L., team, Wellcome Sanger Institute Scientific Operations: Sequencing Operations, Wellcome Sanger Institute Tree of Life Core Informatics team, & Tree of Life Core Informatics collective, D. T. o. L. C. (2025). The genome sequence of a flea beetle, Longitarsus flavicornis (Stephens, 1831) (Coleoptera: Chrysomelidae). *Wellcome Open Res*. https://doi.org/10.12688/wellcomeopenres.25209.1

McCulloch, J., Crowley, L. M., University of Oxford and Wytham Woods Genome Acquisition Lab, Darwin Tree of Life Barcoding collective, Wellcome Sanger Institute Tree of Life Management, S. a. L., team, Wellcome Sanger Institute Scientific Operations: Sequencing Operations, Wellcome Sanger Institute Tree of Life Core Informatics team, & Tree of Life Core Informatics collective, D. T. o. L. C. (2024). The genome sequence of a rove beetle, Lordithon lunulatus (Linnaeus, 1760). *Wellcome Open Res*. https://doi.org/10.12688/wellcomeopenres.22746.1

McCulloch, J., Crowley, L. M., University of Oxford and Wytham Woods Genome Acquisition Lab, Darwin Tree of Life Barcoding collective, Wellcome Sanger Institute Tree of Life Management, S. a. L., team, Wellcome Sanger Institute Scientific Operations: Sequencing Operations, Wellcome Sanger Institute Tree of Life Core Informatics team, & Tree of Life Core Informatics collective, D. T. o. L. C. (2025). The genome sequence of a rove beetle, Ontholestes murinus (Linnaeus, 1758). *Wellcome Open Res*. https://doi.org/10.12688/wellcomeopenres.24330.1

McCulloch, J., University of, O., Wytham Woods Genome Acquisition, L., Darwin Tree of Life Barcoding, c., Wellcome Sanger Institute Tree of Life Management, S., Laboratory, t., Wellcome Sanger Institute Scientific Operations: Sequencing, O., Wellcome Sanger Institute Tree of Life Core Informatics, t., Tree of Life Core Informatics, c., & Darwin Tree of Life, C. (2024). The genome sequence of knotweed leaf beetle, Gastrophysa polygoni (Linnaeus, 1758). *Wellcome Open Res*, *9*, 642. https://doi.org/10.12688/wellcomeopenres.23293.1

McCulloch, J., University of, O., Wytham Woods Genome Acquisition, L., Darwin Tree of Life Barcoding, c., Wellcome Sanger Institute Tree of Life, p., Wellcome Sanger Institute Scientific Operations, D. N. A. P. c., Tree of Life Core Informatics, c., & Darwin Tree of Life, C. (2023). The genome sequence of a rove beetle, Othius punctulatus (Goeze, 1777). *Wellcome Open Res*, *8*, 519. https://doi.org/10.12688/wellcomeopenres.20338.1

McCulloch, J., University of Oxford and Wytham Woods Genome Acquisition Lab, Darwin Tree of Life Barcoding collective, Wellcome Sanger Institute Tree of Life Management, S. a. L., team, Wellcome Sanger Institute Scientific Operations: Sequencing Operations, Wellcome Sanger Institute Tree of Life Core Informatics team, & Tree of Life Core Informatics collective, D. T. o. L. C. (2024). The genome sequence of the greater thorn-tipped longhorn beetle, Pogonocherus hispidulus (Pitter and Mitterpacher, 1783). *Wellcome Open Res*. https://doi.org/10.12688/wellcomeopenres.22455.1

McCulloch, J., University of Oxford and Wytham Woods Genome Acquisition Lab, Darwin Tree of Life Barcoding collective, Wellcome Sanger Institute Tree of Life Management, S. a. L., team, Wellcome Sanger Institute Scientific Operations: Sequencing Operations, Wellcome Sanger Institute Tree of Life Core Informatics team, & Tree of Life Core Informatics collective, D. T. o. L. C. (2025). The genome sequence of a rove beetle, Tachinus rufipes (Linnaeus, 1758). *Wellcome Open Res*. https://doi.org/10.12688/wellcomeopenres.23741.1

Mitchell, R., Geiser, M. F., Barclay, M. V. L., Natural History Museum Genome Acquisition, L., Darwin Tree of Life Barcoding, c., Wellcome Sanger Institute Tree of Life Management, S., Laboratory, t., Wellcome Sanger Institute Scientific Operations: Sequencing, O., Wellcome Sanger Institute Tree of Life Core Informatics, t., Tree of Life Core Informatics, c., & Darwin Tree of Life, C. (2025). The genome sequence of a soldier beetle, Malthinus seriepunctatus Kiesenwetter, 1851. *Wellcome Open Res*, *10*, 80. https://doi.org/10.12688/wellcomeopenres.23716.1

Mitchell, R., Geiser, M. F., Turner, T., Natural History Museum Genome Acquisition, L., Darwin Tree of Life Barcoding, c., Wellcome Sanger Institute Tree of Life Management, S., Laboratory, t., Wellcome Sanger Institute Scientific Operations: Sequencing, O., Wellcome Sanger Institute Tree of Life Core Informatics, t., Tree of Life Core Informatics, c., & Darwin Tree of Life, C. (2025). The genome sequence of the Rock-rose Pot Beetle, Cryptocephalus primarius Harold, 1872. *Wellcome Open Res*, *10*, 77. https://doi.org/10.12688/wellcomeopenres.23703.1

Mitchell, R. C., L. M.; Natural History Museum Genome Acquisition Lab,, University of Oxford and Wytham Woods Genome Acquisition Lab, Darwin Tree of Life Barcoding collective, Wellcome Sanger Institute Tree of Life Management, S. a. L., team, Wellcome Sanger Institute Scientific Operations: Sequencing Operations, Wellcome Sanger Institute Tree of Life Core Informatics team, & Tree of Life Core Informatics collective, D. T. o. L. C. (2024). The genome sequence of the False Ladybird, Endomychus coccineus (Linnaeus, 1758). *Wellcome Open Res*. https://doi.org/10.12688/wellcomeopenres.23097.1

Moran, S., Natural History Museum Genome Acquisition, L., Darwin Tree of Life Barcoding, c., Wellcome Sanger Institute Tree of Life Management, S., Laboratory, t., Wellcome Sanger Institute Scientific Operations: Sequencing, O., Wellcome Sanger Institute Tree of Life Core Informatics, t., Tree of Life Core Informatics, c., & Darwin Tree of Life, C. (2024). The genome sequence of the jumping weevil, Orchestes rusci (Herbst, 1795). *Wellcome Open Res*, *9*, 398. https://doi.org/10.12688/wellcomeopenres.22745.1

Olga Sivell, D. S., Maxwell V. L. Barclay, Liam M. Crowley,, Natural History Museum Genome Acquisition Lab, University of Oxford and Wytham Woods Genome Acquisition Lab, Darwin Tree of Life Barcoding collective, Wellcome Sanger Institute Tree of Life Management, S. a. L., team, Wellcome Sanger Institute Scientific Operations: Sequencing Operations, Wellcome Sanger Institute Tree of Life Core Informatics team, & Tree of Life Core Informatics collective, D. T. o. L. C. (2023). The genome sequence of a longhorn beetle, Rutpela maculata (Poda, 1769). *Wellcome Open Res*. https://doi.org/10.12688/wellcomeopenres.20500.1

Oppert, B., Dossey, A. T., Chu, F. C., Satovic-Vuksic, E., Plohl, M., Smith, T. P. L., Koren, S., Olmstead, M. L., Leierer, D., Ragan, G., & Johnston, J. S. (2023). The Genome of the Yellow Mealworm, Tenebrio molitor: It's Bigger Than You Think. *Genes (Basel)*, *14*(12). https://doi.org/10.3390/genes14122209

Orledge, G. M., Crowley, L. M., University of, O., Wytham Woods Genome Acquisition, L., Darwin Tree of Life Barcoding, c., Wellcome Sanger Institute Tree of Life Management, S., Laboratory, t., Wellcome Sanger Institute Scientific Operations: Sequencing, O., Wellcome Sanger Institute Tree of Life Core Informatics, t., Tree of Life Core Informatics, c., & Darwin Tree of Life, C. (2025). The genome sequence of Eledona agricola (Herbst, 1783) (Coleoptera: Tenebrionidae). *Wellcome Open Res*, *10*, 501. https://doi.org/10.12688/wellcomeopenres.24809.1

Oxford, G. S., Natural History Museum Genome Acquisition, L., Darwin Tree of Life Barcoding, c., Wellcome Sanger Institute Tree of Life Management, S., Laboratory, t., Wellcome Sanger Institute Scientific Operations: Sequencing, O., Wellcome Sanger Institute Tree of Life Core Informatics, t., Tree of Life Core Informatics, c., & Darwin Tree of Life, C. (2025). The genome sequence of the Tansy Beetle, Chrysolina graminis (Linnaeus, 1758). *Wellcome Open Res*, *10*, 296. https://doi.org/10.12688/wellcomeopenres.24287.1

Pang, B., Zhan, Z., & Wang, Y. (2024). A chromosome-level genome assembly of Prosopocoilus inquinatus Westwood, 1848 (Coleoptera: Lucanidae). *Sci Data*, *11*(1), 808. https://doi.org/10.1038/s41597-024-03647-9

Paul, J., Crowley, L. M., University of, O., Wytham Woods Genome Acquisition, L., Darwin Tree of Life Barcoding, c., Wellcome Sanger Institute Tree of Life Management, S., Laboratory, t., Wellcome Sanger Institute Scientific Operations: Sequencing, O., Wellcome Sanger Institute Tree of Life Core Informatics, t., Tree of Life Core Informatics, c., & Darwin Tree of Life, C. (2025). The genome sequence of a flea beetle, Altica lythri Aube, 1843. *Wellcome Open Res*, *10*, 297. https://doi.org/10.12688/wellcomeopenres.24269.1

Paul, J., Crowley, L. M., University of, O., Wytham Woods Genome Acquisition, L., Darwin Tree of Life Barcoding, C., Wellcome Sanger Institute Tree of Life Management, S., Laboratory, t., Wellcome Sanger Institute Scientific Operations: Sequencing, O., Wellcome Sanger Institute Tree of Life Core Informatics, t., Tree of Life Core Informatics, c., & Darwin Tree of Life, C. (2025). The genome sequence of a seed weevil, Oxystoma pomonae (Fabricius, 1798) (Coleoptera: Apionidae). *Wellcome Open Res*, *10*, 640. https://doi.org/10.12688/wellcomeopenres.25092.1

Pu, D. Q., Wu, X. L., Chen, Z. T., Wei, S. J., Cai, P., & Liu, H. L. (2024). Chromosome-level genome assembly of the giant ladybug Megalocaria dilatata. *Sci Data*, *11*(1), 117. https://doi.org/10.1038/s41597-024-02990-1

Ryan Mitchell, R. P., Natural History Museum Genome Acquisition Lab, Darwin Tree of Life Barcoding collective, Wellcome Sanger Institute Tree of Life Management, S. a. L., team, Wellcome Sanger Institute Scientific Operations: Sequencing Operations, Wellcome Sanger Institute Tree of Life Core Informatics team, & Tree of Life Core Informatics collective, D. T. o. L. C. (2024). The genome sequence of the four-banded longhorn beetle, Leptura quadrifasciata Linnaeus, 1758. *Wellcome Open Res*.

https://doi.org/10.12688
Shen, C., Yang, G., Tang, M., Li, X., Zhu, L., Li, W., Jin, L., Deng, P., Zhang, H., Zhai, Q., Wu, G., & Yan, X. (2025). A chromosome-level genome assembly of Mylabris sibirica Fischer von Waldheim, 1823 (Coleoptera, Meloidae). *Sci Data*, *12*(1), 269. https://doi.org/10.1038/s41597-025-04532-9
Sivell, D., Barclay, M. V. L., Mendel, H., Natural History Museum Genome Acquisition, L., Darwin Tree of Life Barcoding, c., Wellcome Sanger Institute Tree of Life Management, S., Laboratory, t., Wellcome Sanger Institute Scientific Operations: Sequencing, O., Wellcome Sanger Institute Tree of Life Core Informatics, t., Tree of Life Core Informatics, c., & Darwin Tree of Life, C. (2024). The genome sequence of a click beetle, Melanotus villosus (Geoffroy in Fourcroy, 1785). *Wellcome Open Res*, *9*, 108. https://doi.org/10.12688/wellcomeopenres.21087.1
Sivell, D., Natural History Museum Genome Acquisition, L., Darwin Tree of Life Barcoding, c., Wellcome Sanger Institute Tree of Life, p., Wellcome Sanger Institute Scientific Operations, D. N. A. P. c., Tree of Life Core Informatics, c., & Darwin Tree of Life, C. (2021). The genome sequence of the red-headed cardinal beetle, Pyrochroa serraticornis (Scopoli, 1763). *Wellcome Open Res*, *6*, 316. https://doi.org/10.12688/wellcomeopenres.17362.1
Sivell, D., Sivell, O., Barclay, M. V. L., Natural History Museum Genome Acquisition Lab, Darwin Tree of Life Barcoding collective, Wellcome Sanger Institute Tree of Life Management, S. a. L., team, Wellcome Sanger Institute Scientific Operations: Sequencing Operations, Wellcome Sanger Institute Tree of Life Core Informatics team, & Tree of Life Core Informatics collective, D. T. o. L. C. (2025). The genome sequence of a dung beetle, Melinopterus prodromus (Brahm, 1790). *Wellcome Open Res*. https://doi.org/10.12688/wellcomeopenres.23799.1
Sivell, D., Sivell, O., Mitchell, R., Natural History Museum Genome Acquisition, L., Darwin Tree of Life Barcoding, c., Wellcome Sanger Institute Tree of Life Management, S., Laboratory, t., Wellcome Sanger Institute Scientific Operations: Sequencing, O., Wellcome Sanger Institute Tree of Life Core Informatics, t., Tree of Life Core Informatics, c., & Darwin Tree of Life, C. (2025). The genome sequence of a carabid beetle, Carabus problematicus Herbst, 1786. *Wellcome Open Res*, *10*, 59. https://doi.org/10.12688/wellcomeopenres.23689.1
Sivell, D., Telnov, D., Geiser, M. F., Barclay, M. V. L., Natural History Museum Genome Acquisition, L., Darwin Tree of Life Barcoding, c., Wellcome Sanger Institute Tree of Life Management, S., Laboratory, t., Wellcome Sanger Institute Scientific Operations: Sequencing, O., Wellcome Sanger Institute Tree of Life Core Informatics, t., Tree of Life Core Informatics, c., & Darwin Tree of Life, C. (2025). The genome sequence of the click beetle, Ampedus sanguinolentus sanguinolentus (Schrank, 1776). *Wellcome Open Res*, *10*, 96. https://doi.org/10.12688/wellcomeopenres.23712.1
Sivell, O., Levey, B., Barclay, M. V. L., Natural History Museum Genome Acquisition, L., Darwin Tree of Life Barcoding, c., Wellcome Sanger Institute Tree of Life, p., Wellcome Sanger Institute Scientific Operations, D. N. A. P. c., Tree of Life Core Informatics, c., & Darwin Tree of Life, C. (2023). The genome sequence of a darkling beetle, Lagria hirta (Linnaeus, 1758). *Wellcome Open Res*, *8*, 501. https://doi.org/10.12688/wellcomeopenres.20232.1
Sivell, O., Sivell, D., Geiser, M., Natural History Museum Genome Acquisition, L., Darwin Tree

of Life Barcoding, c., Wellcome Sanger Institute Tree of Life, p., Wellcome Sanger Institute Scientific Operations, D. N. A. P. c., Tree of Life Core Informatics, c., & Darwin Tree of Life, C. (2023). The genome sequence of a soldier beetle, Cantharis rufa (Linnaeus, 1758). *Wellcome Open Res*, *8*, 478. https://doi.org/10.12688/wellcomeopenres.19986.1

Sivell, O., Sivell, D., Geiser, M., Natural History Museum Genome Acquisition, L., Darwin Tree of Life Barcoding, c., Wellcome Sanger Institute Tree of Life, p., Wellcome Sanger Institute Scientific Operations, D. N. A. P. c., Tree of Life Core Informatics, c., & Darwin Tree of Life, C. (2023). The genome sequence of the Cow Parsley Leaf Beetle, Chrysolina oricalcia (O.F. Muller, 1776). *Wellcome Open Res*, *8*, 400. https://doi.org/10.12688/wellcomeopenres.19985.2

Sivell, O., Sivell, D., Mitchell, R., Barclay, M. V. L., Natural History Museum Genome Acquisition, L., Darwin Tree of Life Barcoding, c., Wellcome Sanger Institute Tree of Life Management, S., Laboratory, t., Wellcome Sanger Institute Scientific Operations: Sequencing, O., Wellcome Sanger Institute Tree of Life Core Informatics, t., Tree of Life Core Informatics, c., & Darwin Tree of Life, C. (2025). The genome sequence of a carabid beetle, Abax parallelepipedus (Piller & Mitterpacher, 1783). *Wellcome Open Res*, *10*, 147. https://doi.org/10.12688/wellcomeopenres.23888.1

Sivell, O., Sivell, D., Mitchell, R., Natural History Museum Genome Acquisition, L., Darwin Tree of Life Barcoding, c., Wellcome Sanger Institute Tree of Life, p., Wellcome Sanger Institute Scientific Operations, D. N. A. P. c., Tree of Life Core Informatics, c., & Darwin Tree of Life, C. (2023). The genome sequence of a leaf beetle, Cryptocephalus moraei (Linnaeus, 1758). *Wellcome Open Res*, *8*, 467. https://doi.org/10.12688/wellcomeopenres.19522.1

Sivell, O., Sivell, D., Natural History Museum Genome Acquisition, L., Darwin Tree of Life Barcoding, c., Wellcome Sanger Institute Tree of Life, p., Wellcome Sanger Institute Scientific Operations, D. N. A. P. c., Tree of Life Core Informatics, c., & Darwin Tree of Life, C. (2023). The genome sequence of a carabid beetle, Nebria salina (Fairmaire & Laboulbene, 1854). *Wellcome Open Res*, *8*, 247. https://doi.org/10.12688/wellcomeopenres.19372.1

Sivell, O., Sivell, D., Natural History Museum Genome Acquisition Lab, Darwin Tree of Life Barcoding collective, Wellcome Sanger Institute Tree of Life programme, Wellcome Sanger Institute Scientific Operations: DNA Pipelines collective, & Tree of Life Core Informatics collective, D. T. o. L. C. (2021). The genome sequence of a soldier beetle, Cantharis rustica (Fallén 1807). *Wellcome Open Res*. https://doi.org/10.12688/wellcomeopenres.17363.1

Sivell, O., Taylor, S. C., Barclay, M. V. L., Skipp, S., Geiser, M. F., Natural History Museum Genome Acquisition, L., Darwin Tree of Life Barcoding, c., Wellcome Sanger Institute Tree of Life Management, S., Laboratory, t., Wellcome Sanger Institute Scientific Operations: Sequencing, O., Wellcome Sanger Institute Tree of Life Core Informatics, t., Tree of Life Core Informatics, c., & Darwin Tree of Life, C. (2025). The genome sequence of a beetle, Pycnomerus fuliginosus Erichson, 1842. *Wellcome Open Res*, *10*, 144. https://doi.org/10.12688/wellcomeopenres.23770.1

Sivell, O. L., B.; Barclay, M. V.L.; Sivell, D.; Natural History Museum Genome Acquisition Lab,, Darwin Tree of Life Barcoding collective, Wellcome Sanger Institute Tree of Life Management, S. a. L., team, Wellcome Sanger Institute Scientific Operations:

Sequencing Operations, Wellcome Sanger Institute Tree of Life Core Informatics team, & Tree of Life Core Informatics collective, D. T. o. L. C. (2025). The genome sequence of a weevil, Philopedon plagiatum (Schaller, 1783). *Wellcome Open Res*. https://doi.org/10.12688/wellcomeopenres.23784.1

Sladecek, F., Crowley, L. M., University of, O., Wytham Woods Genome Acquisition, L., Darwin Tree of Life Barcoding, c., Wellcome Sanger Institute Tree of Life Management, S., Laboratory, t., Wellcome Sanger Institute Scientific Operations: Sequencing, O., Wellcome Sanger Institute Tree of Life Core Informatics, t., Tree of Life Core Informatics, c., & Darwin Tree of Life, C. (2025). The genome sequence of a small dung beetle, Volinus sticticus (Panzer, 1798), formerly known as Aphodius sticticus. *Wellcome Open Res*, *10*, 195. https://doi.org/10.12688/wellcomeopenres.23692.1

Sladecek, F., Lewis, O. T., University of, O., Wytham Woods Genome Acquisition, L., Darwin Tree of Life Barcoding, c., Wellcome Sanger Institute Tree of Life Management, S., Laboratory, t., Wellcome Sanger Institute Scientific Operations: Sequencing, O., Wellcome Sanger Institute Tree of Life Core Informatics, t., Tree of Life Core Informatics, c., & Darwin Tree of Life, C. (2025). The genome sequence of a dung beetle, Geotrupes spiniger (Marsham, 1802). *Wellcome Open Res*, *10*, 71. https://doi.org/10.12688/wellcomeopenres.23705.2

Spilling, C. S., O.; Kusy, D.; Natural History Museum Genome Acquisition Lab,, Darwin Tree of Life Barcoding collective, Wellcome Sanger Institute Tree of Life Management, S. a. L., team, Wellcome Sanger Institute Scientific Operations: Sequencing Operations, Wellcome Sanger Institute Tree of Life Core Informatics team, & Tree of Life Core Informatics collective, D. T. o. L. C. (2024). The genome sequence of the Orchid Beetle, Dascillus cervinus (Linnaeus, 1758). *Wellcome Open Res*. https://doi.org/10.12688/wellcomeopenres.21161.1

Stahlke, A. R., Ozsoy, A. Z., Bean, D. W., & Hohenlohe, P. A. (2019). Mitochondrial Genome Sequences of Diorhabda carinata and Diorhabda carinulata, Two Beetle Species Introduced to North America for Biological Control. *Microbiol Resour Announc*, *8*(35). https://doi.org/10.1128/MRA.00690-19

Sylvester, T., Hoover, Z., Hjelmen, C. E., Jonika, M. M., Blackmon, L. T., Alfieri, J. M., Johnston, J. S., Chien, S., Esfandani, T., & Blackmon, H. (2024). A reference quality genome assembly for the jewel scarab Chrysina gloriosa. *G3 (Bethesda)*, *14*(6). https://doi.org/10.1093/g3journal/jkae084

Talay Namintraporn, M. V. L. B., Michael F. Geiser,, Keita Matsumoto, D. V., Joana Cristóvão, Will Bayfield-Farrell ,, Ian Sims, N. H. M. G. A. L., Darwin Tree of Life Barcoding collective, Wellcome Sanger Institute Tree of Life Management, S. a. L., team, Wellcome Sanger Institute Scientific Operations: Sequencing Operations, Wellcome Sanger Institute Tree of Life Core Informatics team, & Tree of Life Core Informatics collective, D. T. o. L. C. (2025). The genome sequence of the Cream-spot ladybird, Calvia quattuordecimguttata (Linnaeus, 1758). *Wellcome Open Res*. https://doi.org/10.12688/wellcomeopenres.24294.1

Tang, B., Yin, C., He, K., Tao, S., Fu, L., Liu, Y., Li, F., & Hou, Y. (2024). A chromosome-scale genome assembly of the nipa palm hispid beetle Octodonta nipae. *Sci Data*, *11*(1), 562. https://doi.org/10.1038/s41597-024-03417-7

Telfer, M., Booth, R., Phillips, D., University of Oxford and Wytham Woods Genome Acquisition Lab, Natural History Museum Genome Acquisition Lab, Darwin Tree of Life

Barcoding collective, Wellcome Sanger Institute Tree of Life Management, S. a. L., team, Wellcome Sanger Institute Scientific Operations: Sequencing Operations, Wellcome Sanger Institute Tree of Life Core Informatics team, & Tree of Life Core Informatics collective, D. T. o. L. C. (2024). The genome sequence of the narrow-waisted bark beetle, Salpingus planirostris (Fabricius, 1787). *Wellcome Open Res*. https://doi.org/10.12688/wellcomeopenres.22985.1

Telfer, M., Crowley, L., Barclay, M., Bickerstaff, J., University of Oxford and Wytham Woods Genome Acquisition Lab, Natural History Museum Genome Acquisition Lab, Darwin Tree of Life Barcoding collective, Wellcome Sanger Institute Tree of Life Management, S. a. L., team, Wellcome Sanger Institute Scientific Operations: Sequencing Operations, Wellcome Sanger Institute Tree of Life Core Informatics team, & Tree of Life Core Informatics collective, D. T. o. L. C. (2024). The genome sequence of the hairy spider weevil, Barypeithes pellucidus (Boheman, 1834). *Wellcome Open Res*. https://doi.org/10.12688/wellcomeopenres.23067.1

Telfer, M. G., Badham, X. R., University of, O., Wytham Woods Genome Acquisition, L., Darwin Tree of Life Barcoding, c., Wellcome Sanger Institute Tree of Life Management, S., Laboratory, t., Wellcome Sanger Institute Scientific Operations: Sequencing, O., Wellcome Sanger Institute Tree of Life Core Informatics, t., Tree of Life Core Informatics, c., & Darwin Tree of Life, C. (2024). The genome sequence of the beech bark beetle, Taphrorychus bicolor (Herbst, 1793). *Wellcome Open Res*, *9*, 213. https://doi.org/10.12688/wellcomeopenres.21265.1

Telfer, M. G., Barclay, M. V. L., Phillips, D., University of, O., Wytham Woods Genome Acquisition, L., Natural History Museum Genome Acquisition, L., Darwin Tree of Life Barcoding, c., Wellcome Sanger Institute Tree of Life Management, S., Laboratory, t., Wellcome Sanger Institute Scientific Operations: Sequencing, O., Wellcome Sanger Institute Tree of Life Core Informatics, t., Tree of Life Core Informatics, c., & Darwin Tree of Life, C. (2024). The genome sequence of a false flower beetle, Anaspis maculata (Geoffroy in Fourcroy, 1785). *Wellcome Open Res*, *9*, 212. https://doi.org/10.12688/wellcomeopenres.21283.1

Telfer, M. G., Bickerstaff, J., University of, O., Wytham Woods Genome Acquisition, L., Darwin Tree of Life Barcoding, c., Wellcome Sanger Institute Tree of Life Management, S., Laboratory, t., Wellcome Sanger Institute Scientific Operations: Sequencing, O., Wellcome Sanger Institute Tree of Life Core Informatics, t., Tree of Life Core Informatics, c., & Darwin Tree of Life, C. (2024). The genome sequence of an Entiminae weevil, Polydrusus pterygomalis Boheman, 1840. *Wellcome Open Res*, *9*, 528. https://doi.org/10.12688/wellcomeopenres.23048.1

Telfer, M. G., Blomfield-Smith, H., University of, O., Wytham Woods Genome Acquisition, L., Darwin Tree of Life Barcoding, c., Wellcome Sanger Institute Tree of Life Management, S., Laboratory, t., Wellcome Sanger Institute Scientific Operations: Sequencing, O., Wellcome Sanger Institute Tree of Life Core Informatics, t., Tree of Life Core Informatics, c., & Darwin Tree of Life, C. (2024). The genome sequence of the flea beetle, Crepidodera aurea (Geoffrey, 1785). *Wellcome Open Res*, *9*, 318. https://doi.org/10.12688/wellcomeopenres.22454.1

Telfer, M. G., Crowley, L. M., Badham, X. R., University of Oxford and Wytham Woods Genome Acquisition Lab, Darwin Tree of Life Barcoding collective, Wellcome Sanger Institute Tree of Life Management, S. a. L., team, Wellcome Sanger Institute Scientific

Operations: Sequencing Operations, Wellcome Sanger Institute Tree of Life Core Informatics team, & Tree of Life Core Informatics collective, D. T. o. L. C. (2025). The genome sequence of a ground beetle, Poecilus cupreus (Linnaeus, 1758). *Wellcome Open Res*. https://doi.org/10.12688/wellcomeopenres.24263.1

Telfer, M. G., Geiser, M. F., University of, O., Wytham Woods Genome Acquisition, L., Darwin Tree of Life Barcoding, c., Wellcome Sanger Institute Tree of Life Management, S., Laboratory, t., Wellcome Sanger Institute Scientific Operations: Sequencing, O., Wellcome Sanger Institute Tree of Life Core Informatics, t., Tree of Life Core Informatics, c., & Darwin Tree of Life, C. (2024). The genome sequence of a soldier beetle, Malthinus flaveolus (Herbst, 1786). *Wellcome Open Res*, *9*, 121. https://doi.org/10.12688/wellcomeopenres.21086.1

Telfer, M. G., Phillips, D., University of, O., Wytham Woods Genome Acquisition, L., Darwin Tree of Life Barcoding, c., Wellcome Sanger Institute Tree of Life Management, S., Laboratory, t., Wellcome Sanger Institute Scientific Operations: Sequencing, O., Wellcome Sanger Institute Tree of Life Core Informatics, t., Tree of Life Core Informatics, c., & Darwin Tree of Life, C. (2024). The genome sequence of a metallic wood-boring beetle, Agrilus cyanescens (Ratzeburg, 1837). *Wellcome Open Res*, *9*, 46. https://doi.org/10.12688/wellcomeopenres.20877.2

Van Dam, M. H., Cabras, A. A., Henderson, J. B., Rominger, A. J., Perez Estrada, C., Omer, A. D., Dudchenko, O., Lieberman Aiden, E., & Lam, A. W. (2021). The Easter Egg Weevil (Pachyrhynchus) genome reveals syntenic patterns in Coleoptera across 200 million years of evolution. *PLoS Genet*, *17*(8), e1009745. https://doi.org/10.1371/journal.pgen.1009745

Vassiliades, D., Geiser, M. F., Barclay, M. V. L., Bayfield Farrell, W., Cristovao, J., Matsumoto, K., Crowley, L. M., Natural History Museum Genome Acquisition, L., University of, O., Wytham Woods Genome Acquisition, L., Darwin Tree of Life Barcoding, c., Wellcome Sanger Institute Tree of Life Management, S., Laboratory, t., Wellcome Sanger Institute Scientific Operations: Sequencing, O., Wellcome Sanger Institute Tree of Life Core Informatics, t., Tree of Life Core Informatics, c., & Darwin Tree of Life, C. (2025). The genome sequence of the Alder leaf beetle, Agelastica alni (Linnaeus, 1758). *Wellcome Open Res*, *10*, 263. https://doi.org/10.12688/wellcomeopenres.24286.1

Volaric, M., Despot-Slade, E., Veseljak, D., Mestrovic, N., & Mravinac, B. (2022). Reference-Guided De Novo Genome Assembly of the Flour Beetle Tribolium freemani. *Int J Mol Sci*, *23*(11). https://doi.org/10.3390/ijms23115869

Wang, Q., Liu, L., Zhang, S., Wu, H., & Huang, J. (2022). A chromosome-level genome assembly and intestinal transcriptome of Trypoxylus dichotomus (Coleoptera: Scarabaeidae) to understand its lignocellulose digestion ability. *Gigascience*, *11*. https://doi.org/10.1093/gigascience/giac059

Wang, Y. T., Zhang, Y., Ma, C., Ma, W. H., Cao, L. J., Chen, J. C., Song, W., Yang, J. F., Gao, X. Y., Chen, H. S., Tian, Z. Y., Desneux, N., Wei, S. J., & Zhou, Z. S. (2024). Chromosome-level genome assembly of an oligophagous leaf beetle Ophraella communa (Coleoptera: Chrysomelidae). *Sci Data*, *11*(1), 735. https://doi.org/10.1038/s41597-024-03486-8

Wang, Z., Liu, Y., Wang, H., Roy, A., Liu, H., Han, F., Zhang, X., & Lu, Q. (2023). Genome and transcriptome of Ips nitidus provide insights into high-altitude hypoxia adaptation and symbiosis. *iScience*, *26*(10), 107793. https://doi.org/10.1016/j.isci.2023.107793

Whiffin, A., Darwin Tree of Life Barcoding, C., Wellcome Sanger Institute Tree of Life

Management, S., Laboratory, t., Wellcome Sanger Institute Scientific Operations: Sequencing, O., Wellcome Sanger Institute Tree of Life Core Informatics, t., Tree of Life Core Informatics, c., & Darwin Tree of Life, C. (2025). The genome sequence of the larch ladybird beetle, Aphidecta obliterata (Linnaeus, 1758) (Coleoptera: Coccinellidae). *Wellcome Open Res*, *10*, 646. https://doi.org/10.12688/wellcomeopenres.25090.1

Xiao, H., Ma, C., Li, S., Gao, H., Tao, S., & Yin, C. (2024). A chromosome-scale genome assembly of the peanut beetle, Ulomoides dermestoides (Coleptera: Tenebrionidae). *Sci Data*, *11*(1), 1148. https://doi.org/10.1038/s41597-024-04000-w

Xie, W. W., Sheng, L. J., Wan, Y., Weng, X. Q., Liang, G. H., Zhang, F. P., & Chen, H. (2020). The complete mitochondrial genome of Plagiodera versicolora (Laicharting)(Coleoptera: Chrysomelidae). *Mitochondrial DNA B Resour*, *5*(3), 3600-3601. https://doi.org/10.1080/23802359.2020.1829138

Xu, S., Cheng, L., Hu, W., Gao, Y., Cheng, Q., & Lv, J. (2025). The Complete Mitochondrial Genome of the Pest Beetle Latheticus oryzae: Assembly, Annotation, and Phylogenetic Analysis. *Ecol Evol*, *15*(12), e72717. https://doi.org/10.1002/ece3.72717

Yan, J., Zhang, C., Zhang, M., Zhou, H., Zuo, Z., Ding, X., Zhang, R., Li, F., & Gao, Y. (2023). Chromosome-level genome assembly of the Colorado potato beetle, Leptinotarsa decemlineata. *Sci Data*, *10*(1), 36. https://doi.org/10.1038/s41597-023-01950-5

Ye, M., Xie, Y., Jin, J., Huang, C., Ning, K., Liu, Z., Li, H., & Wang, X. (2024). A chromosome-level genome assembly of Serangium japonicum Chapin, 1940 (Coleoptera: Coccinellidae). *Sci Data*, *11*(1), 1421. https://doi.org/10.1038/s41597-024-04197-w

Zhang, L., Li, S., Luo, J., Du, P., Wu, L., Li, Y., Zhu, X., Wang, L., Zhang, S., & Cui, J. (2020). Chromosome-level genome assembly of the predator Propylea japonica to understand its tolerance to insecticides and high temperatures. *Mol Ecol Resour*, *20*(1), 292-307. https://doi.org/10.1111/1755-0998.13100

Zhang, X., Huang, R., Chen, Y., Li, W., Zhang, X., Yang, J., & Lv, J. (2025). Complete mitochondrial genome of Tribolium castaneum (Coleoptera: Tenebrionidae) reared on sauce-flavor Daqu. *Front Insect Sci*, *5*, 1621855. https://doi.org/10.3389/finsc.2025.1621855

Zhu, M., Zhang, J., Yan, J., & Han, Y. (2025). A chromosomal-level genome assembly of Kibakoganea sinica, Bouchard, 2005 (Coleoptera: Scarabaeidae). *Sci Data*, *12*(1), 1012. https://doi.org/10.1038/s41597-025-05347-4
