## Supplemental Figures for "Species-rich and genomically diverse: comparative genomics reveals how fusions, fissions, and sex chromosomes have shaped beetle evolution"

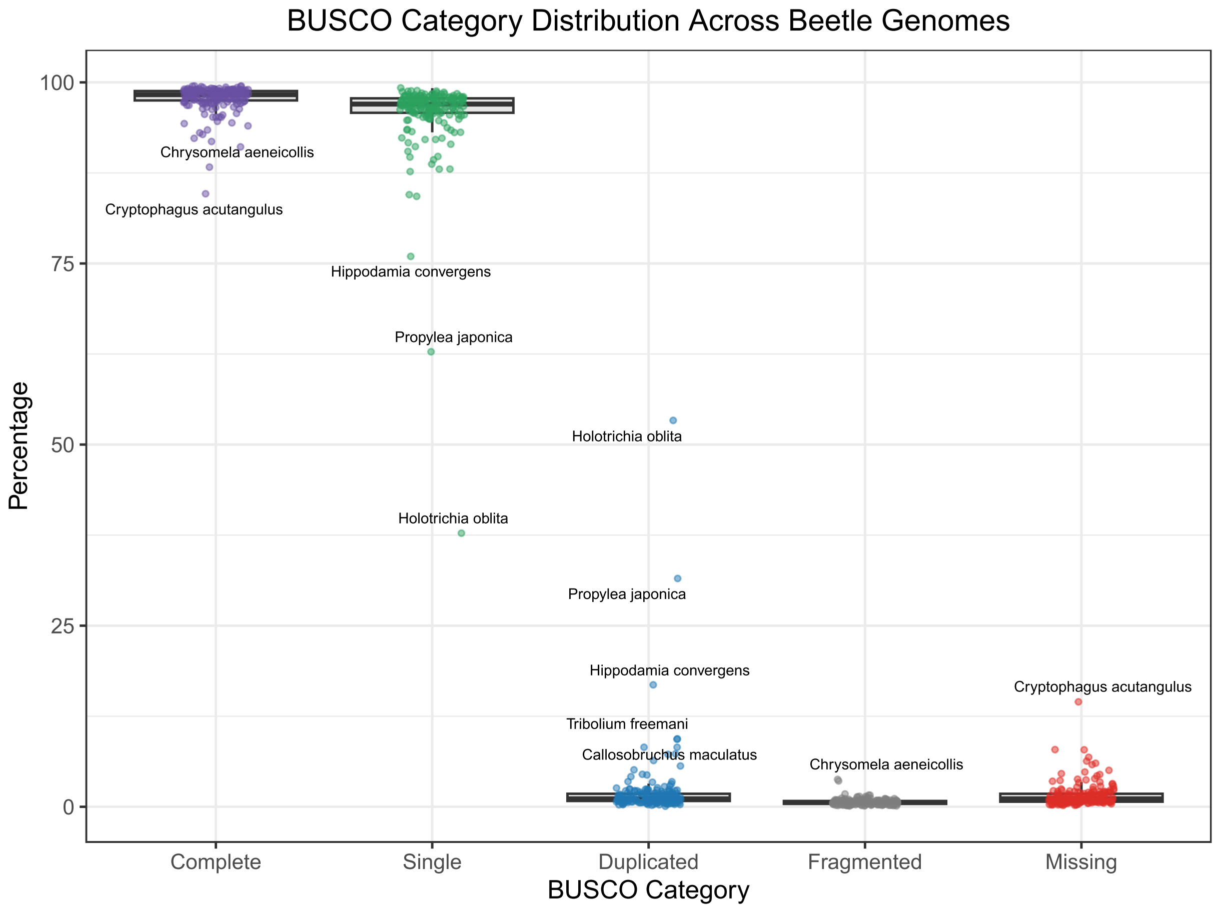


**Supplemental Figure 1. Genome completeness assessment (BUSCO) across beetle species.** Boxplots summarizing the distribution of several BUSCO categories (Complete, Single-copy, Duplicated, Fragmented, and Missing) across 190 analyzed genomes. Points represent individual species with several outliers highlighted.

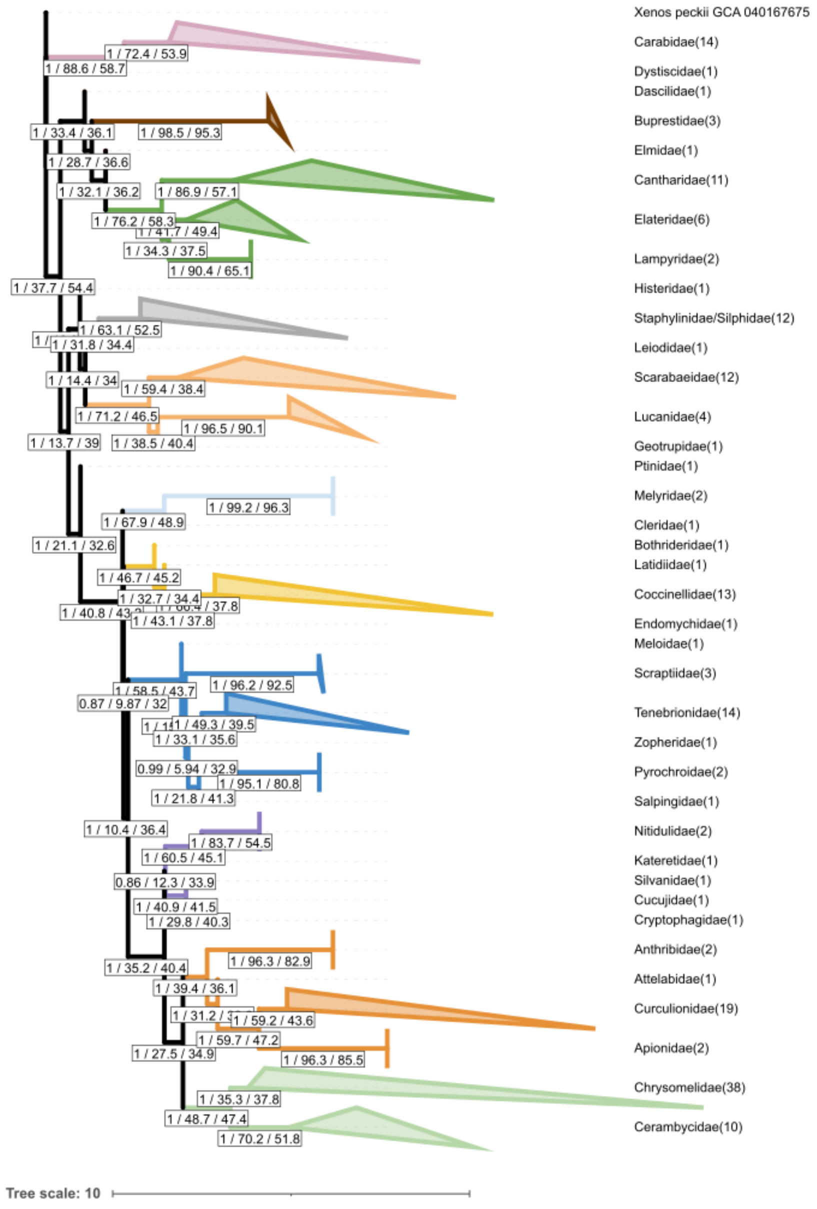


**Supplemental Figure 2. Phylogenetic relationships of beetle families with branch lengths shown.** Families with more than two species were collapsed with the number of total species per family in parentheses. Bootstrap support, gCF and sCF are shown in boxes. Superfamilies are shown color-coded as in Figure 1

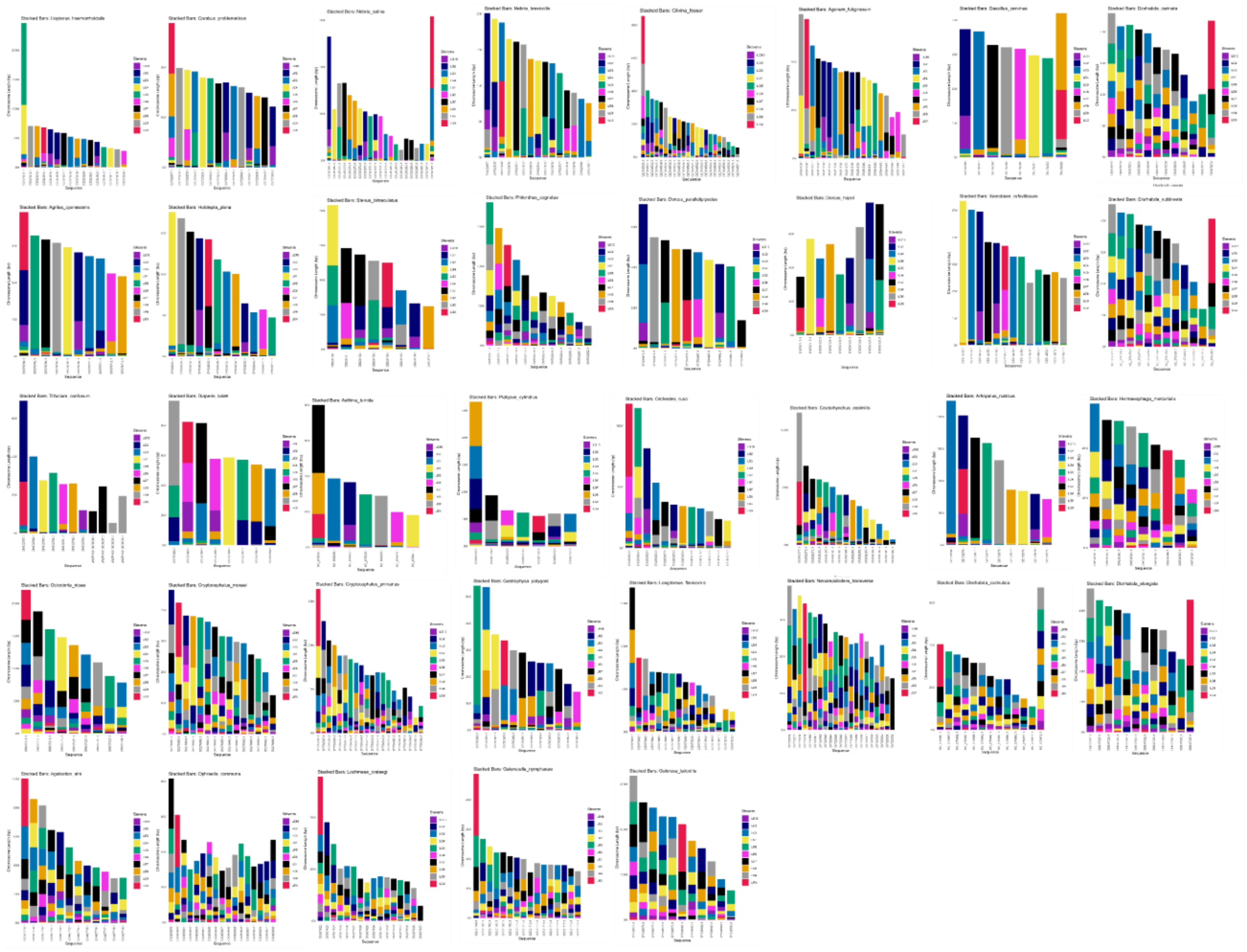

**Supplemental Figure 3. Element_vis plots of Stevens elements across 37 inferred neo-sex chromosome species**. Stacked bar plots represent each chromosome by physical length and the proportion of BUSCOs belonging to particular Stevens elements. Note that Stevens X is shown in red.

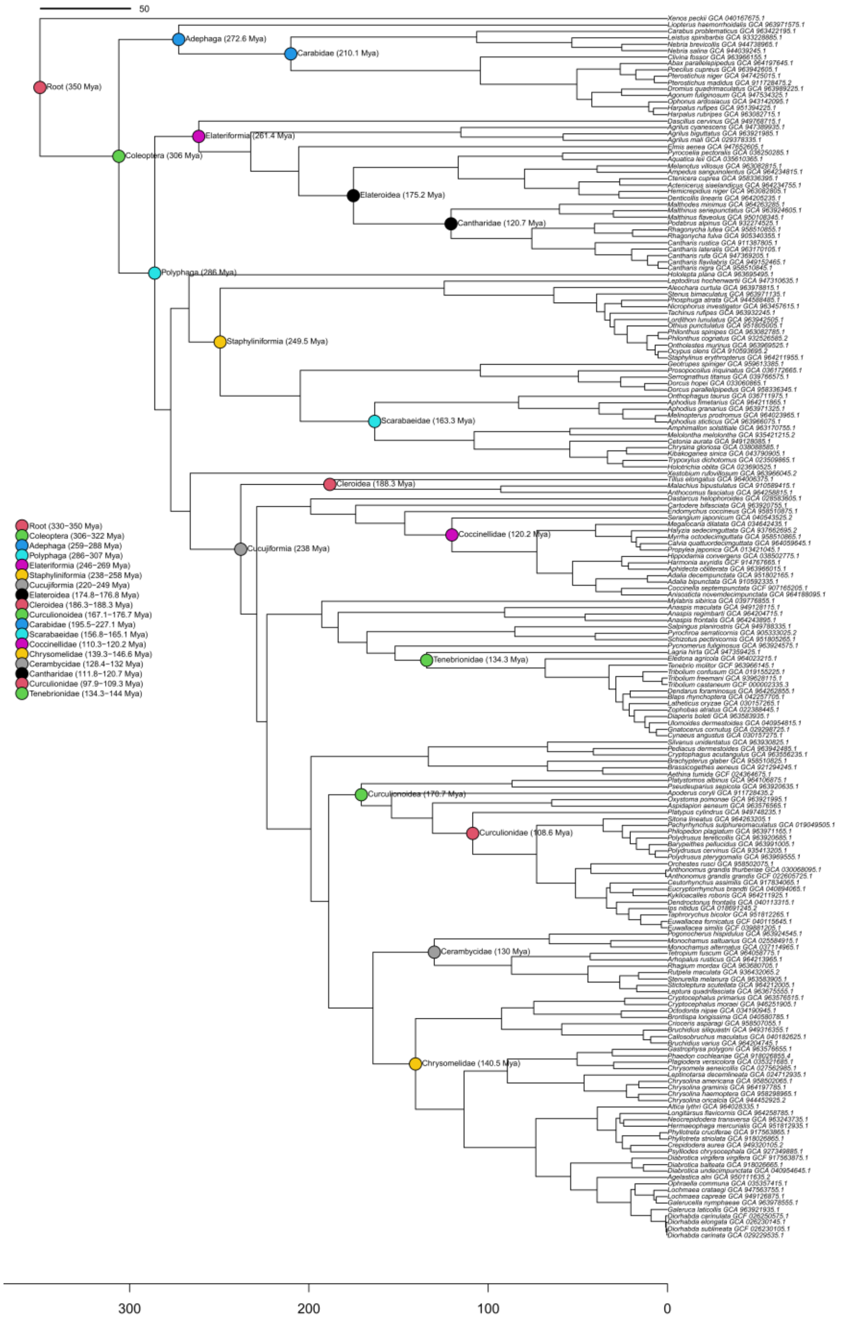


**Supplemental Figure 4. Ultrametric time calibrated tree.** Time-calibrated phylogeny of beetles using time dates from Cai et al. (2022). The ultrametric tree shows the evolutionary relationships with branch lengths proportional to time (millions of years ago). Colored circles indicate divergence times distinct to beetle suborders, infraorder, and family as shown in the legend.

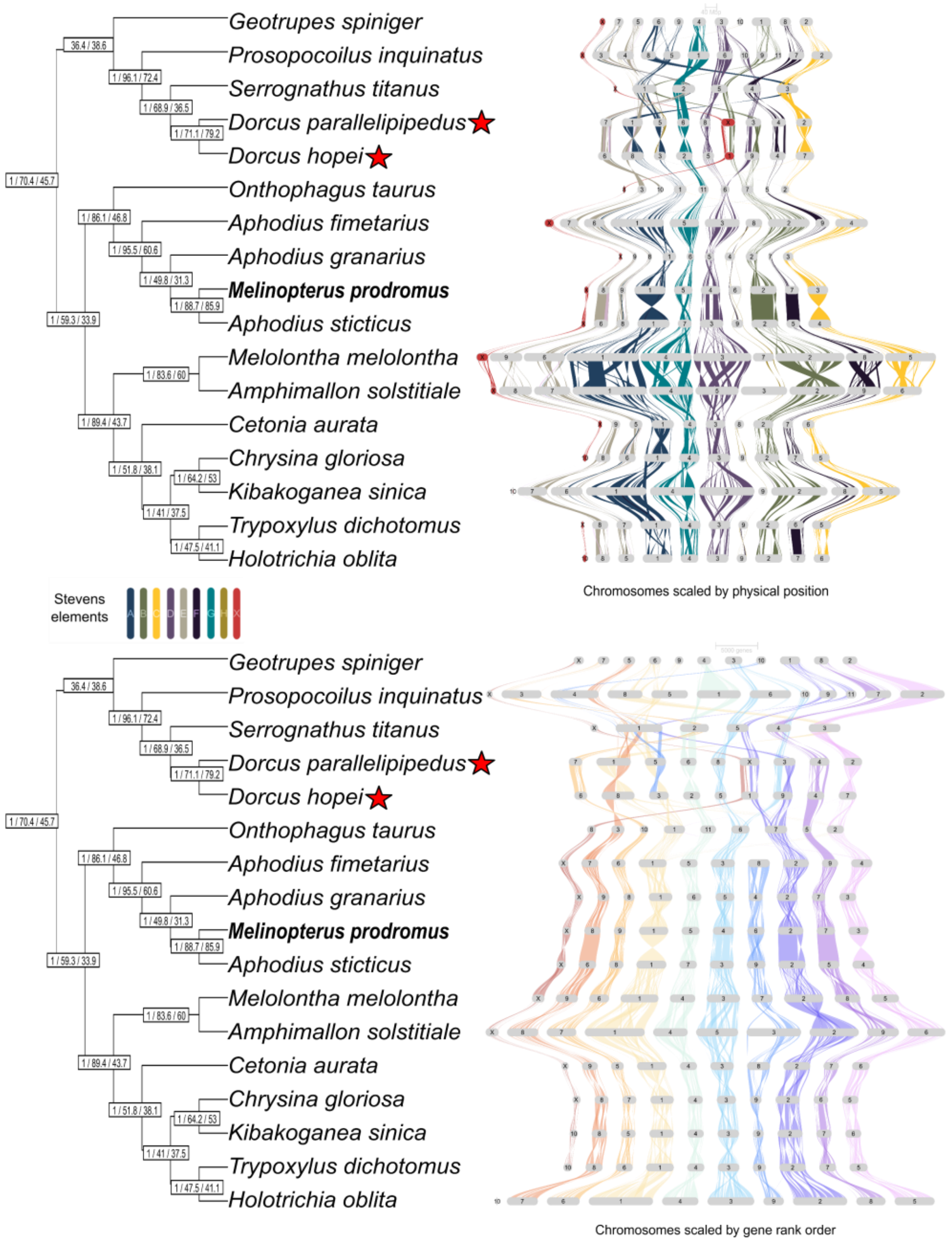


**Supplemental Figure 5. Synteny and collinearity in Scarabaeoidea.** Evolutionary relationships among species in the Scarabaeoidea superfamily with bootstrap support and gCF and sCF shown in boxes. Top panel: GENESPACE plot showing chromosomes scaled by physical position using *Tribolium castaneum* (Stevens elements) as the reference. X chromosome shown in red and labeled X when identified as such in the reference. Bottom panel: GENESPACE plot using gene rank order, and reference *Melinopterus prodromus* (bold on the phylogenetic tree). Red stars highlight species with inferred neo-sex chromosomes.

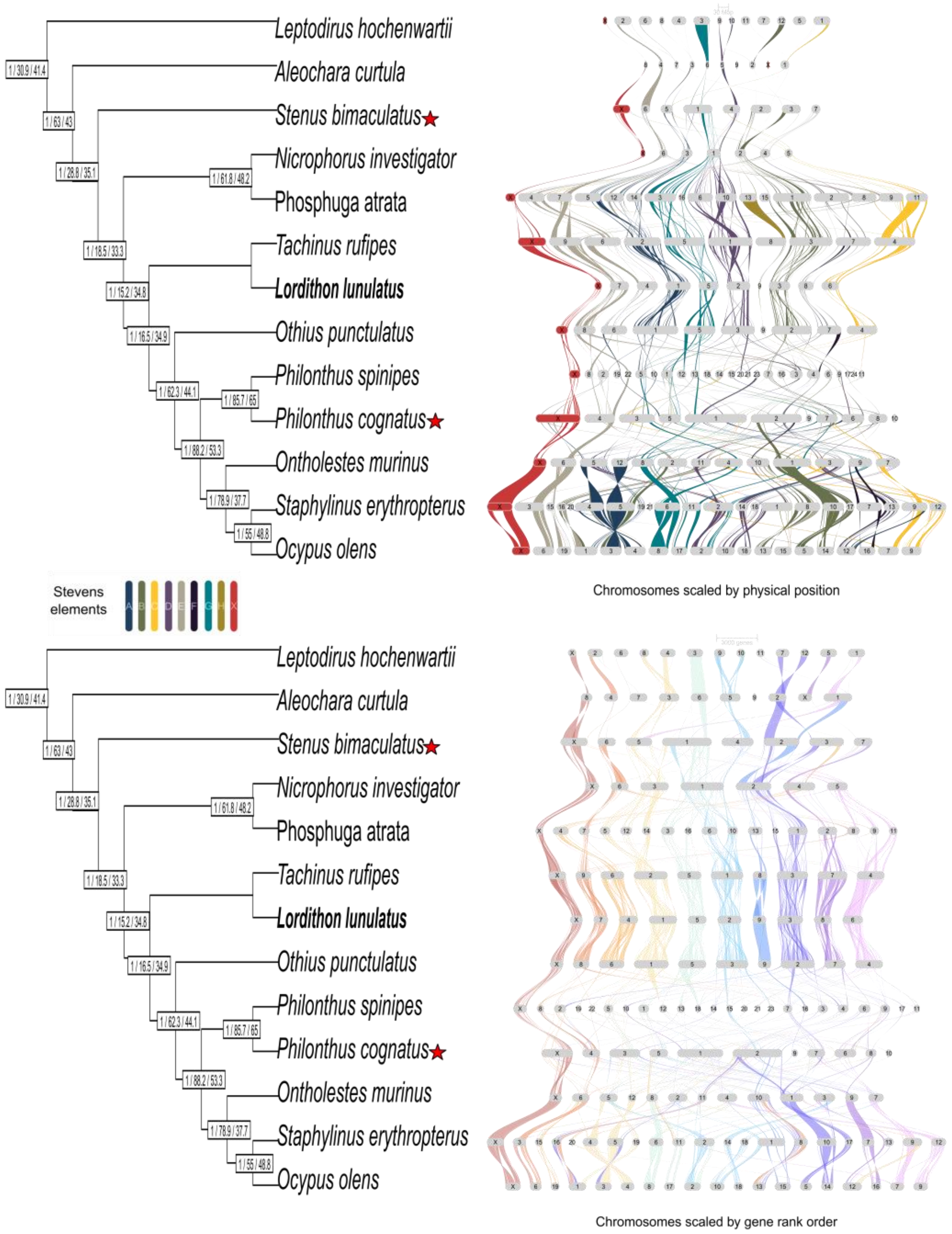


**Supplemental Figure 6. Synteny and collinearity in Staphylinoidea.** Evolutionary relationships among species in the Staphylinoidea superfamily with bootstrap support and gCF and sCF shown in boxes. Top panel: GENESPACE plot showing chromosomes scaled by physical position using *Tribolium castaneum* (Stevens elements) as the reference. X chromosome shown in red and labeled X when identified as such in the reference. Bottom panel: GENESPACE plot using gene rank order, and reference *Lordithon lunulatus* (bold on the phylogenetic tree). Red stars highlight species with inferred neo-sex chromosomes.

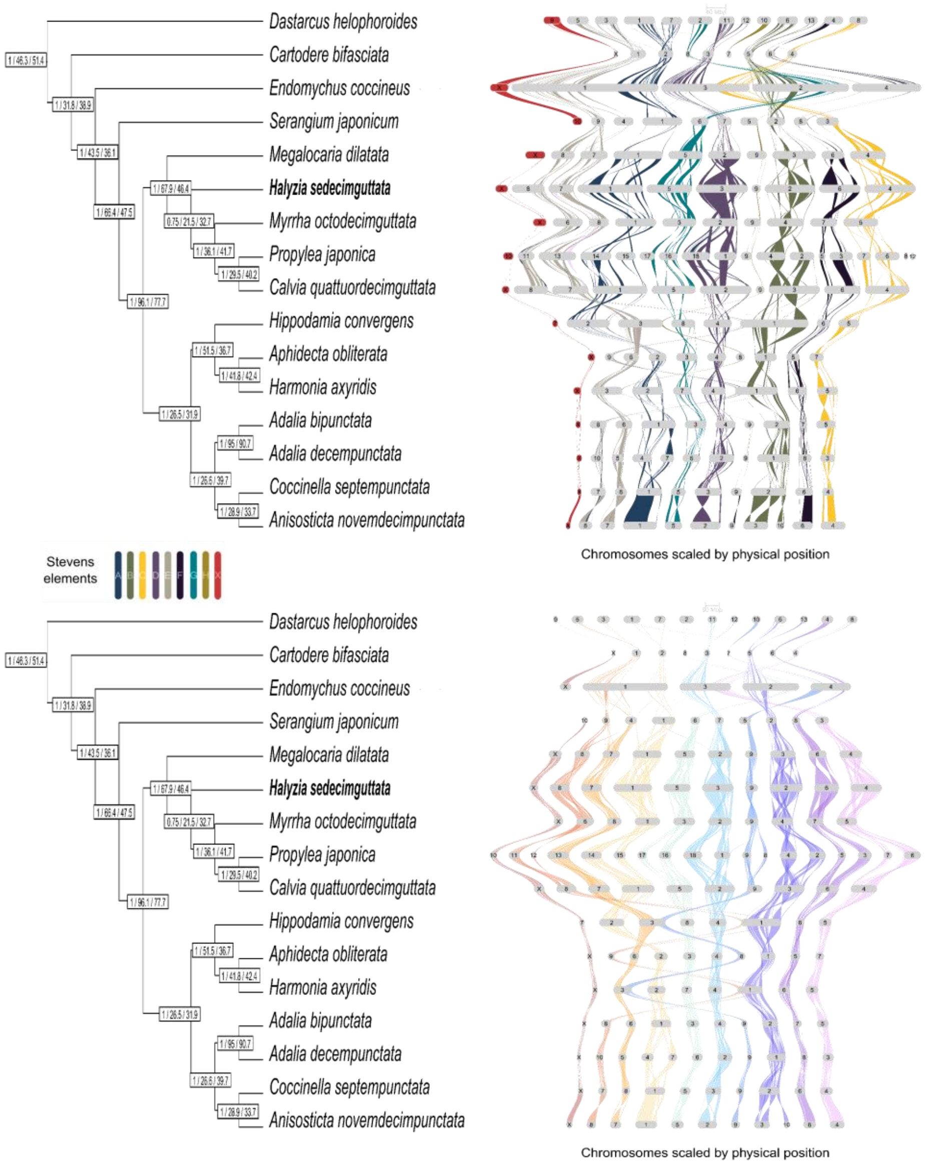


**Supplemental Figure 7. Synteny and collinearity in Coccinelloidea.** Evolutionary relationships among species in the Coccinelloidea superfamily with bootstrap support and gCF and sCF shown in boxes. Top panel: GENESPACE plot showing chromosomes scaled by physical position using *Tribolium castaneum* (Stevens elements) as the reference. X chromosome shown in red and labeled X when identified as such in the reference. Bottom panel: GENESPACE plot using gene rank order, and reference *Halyzia sedecimguttata* (bold on the phylogenetic tree). Red stars highlight species with inferred neo-sex chromosomes.

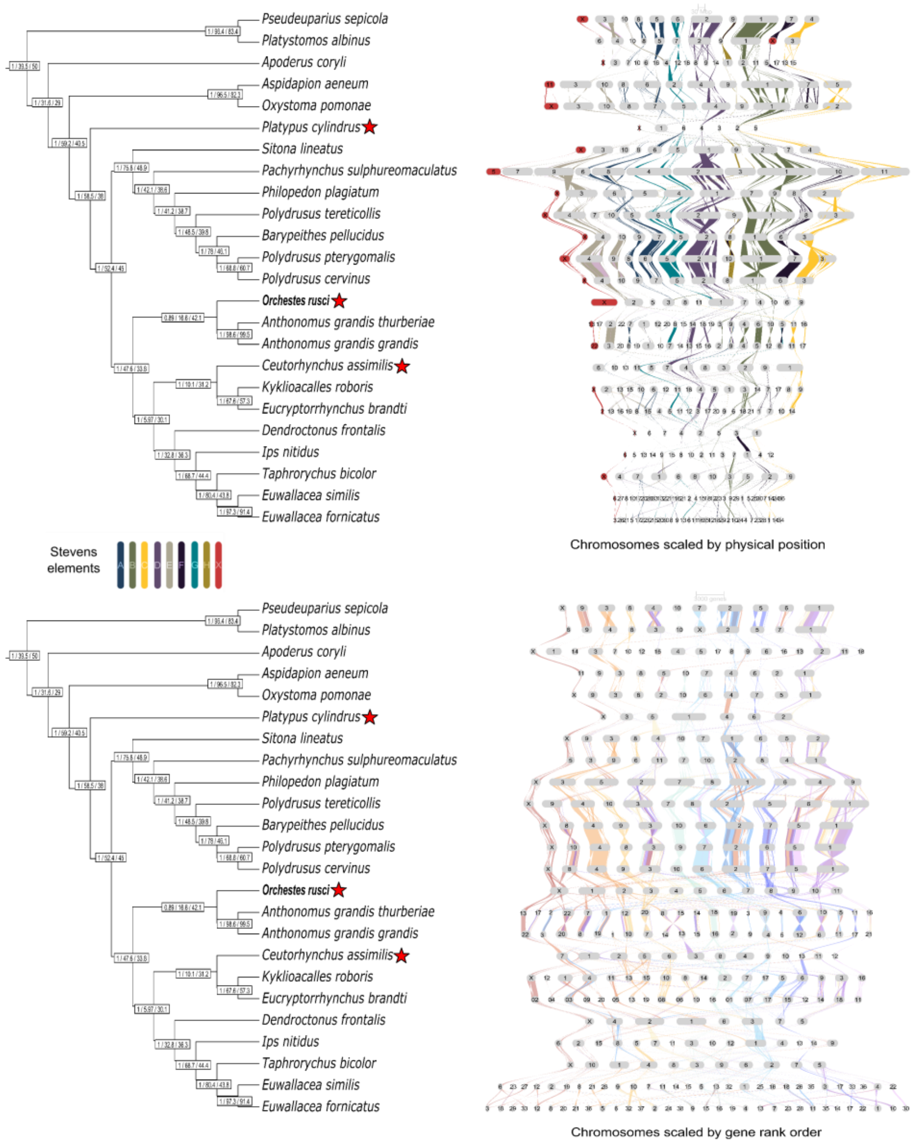


**Supplemental Figure 8. Synteny and collinearity in Curculionoidea.** Evolutionary relationships among species in the Curculionoidea superfamily with bootstrap support and gCF and sCF shown in boxes. Top panel: GENESPACE plot showing chromosomes scaled by physical position using *Tribolium castaneum* (Stevens elements) as the reference. X chromosome shown in red and labeled X when identified as such in the reference. Bottom panel: GENESPACE plot using gene rank order, and reference *Orchestes rusci* (bold on the phylogenetic tree). Red stars highlight species with inferred neo-sex chromosomes.

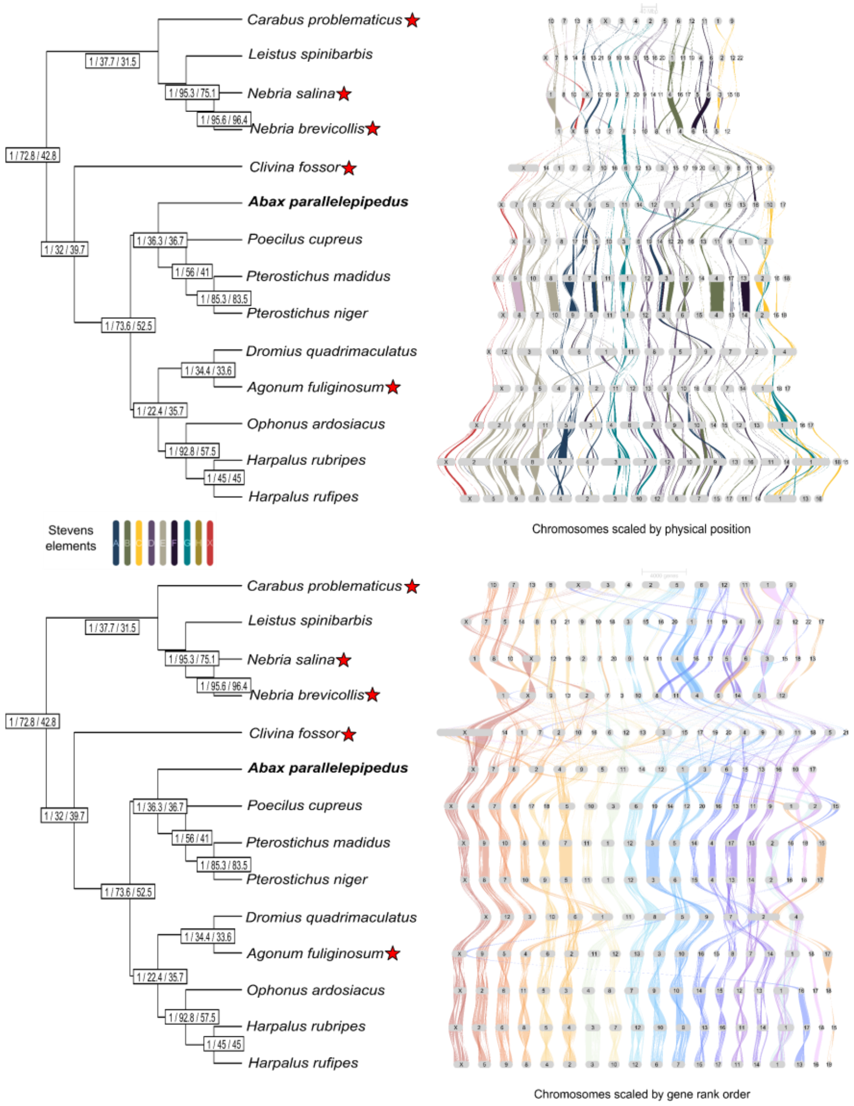


**Supplemental Figure 9. Synteny and collinearity in Caraboidea.** Evolutionary relationships among species in the Curculionoidea superfamily with bootstrap support and gCF and sCF shown in boxes. Top panel: GENESPACE plot showing chromosomes scaled by physical position using *Tribolium castaneum* (Stevens elements) as the reference. X chromosome shown in red and labeled X when identified as such in the reference. Bottom panel: GENESPACE plot using gene rank order, and reference *Abax parallelepipedus* (bold on the phylogenetic tree). Red stars highlight species with inferred neo-sex chromosomes.

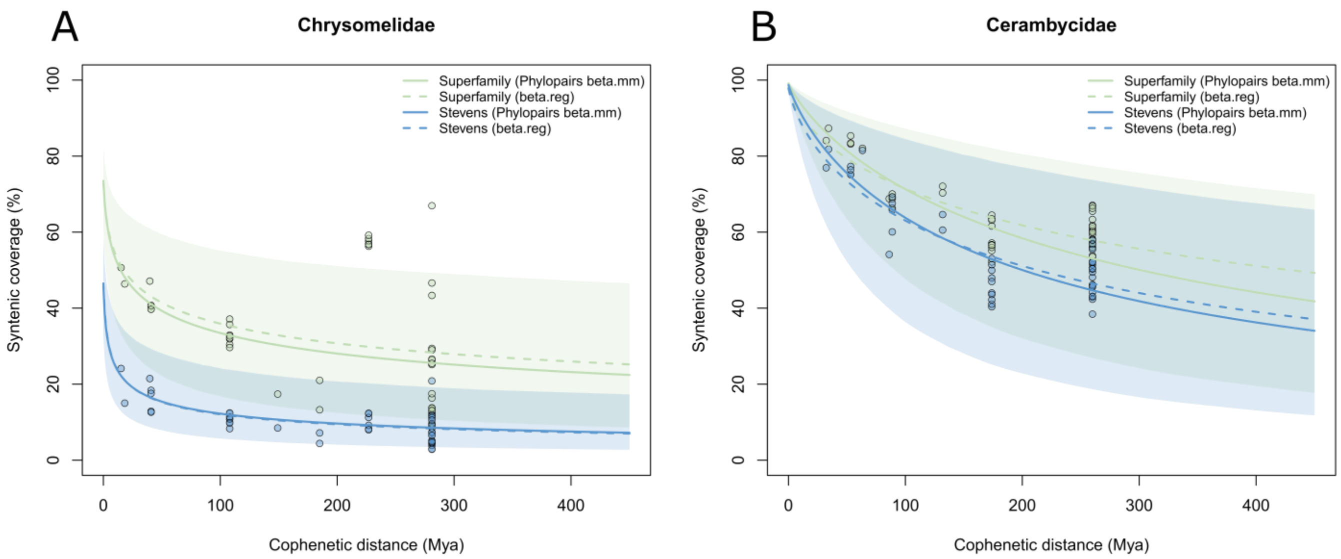


**Supplemental Figure 10. Synteny patterns of the Chrysomelideae vs Cerambycidae.** Plots are similar to **Fig. 5**, yet are broken down here to family level to highlight differences between these two groups. **A)** shows pairwise relationships of 10 randomly sampled Chyrsomelidae, which was equivalent to the total number of **B)** Cerambycidae available for analysis.

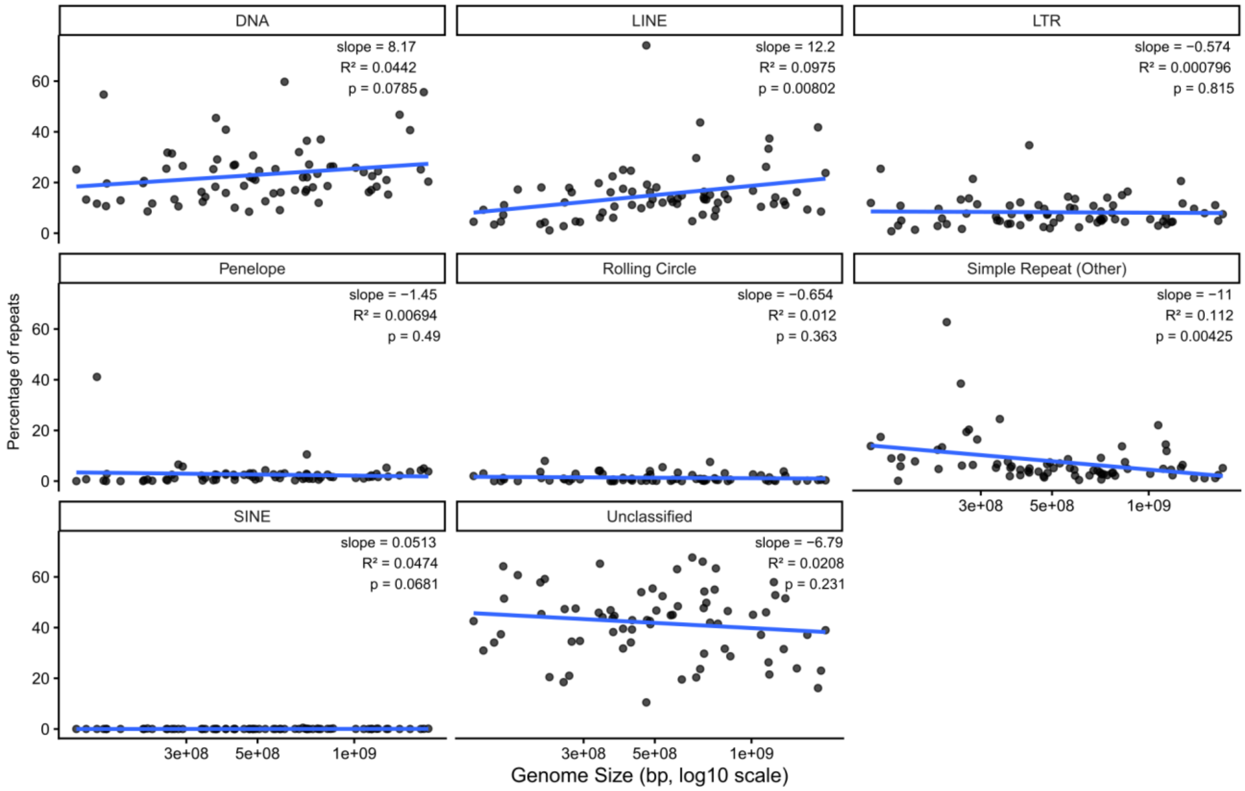


**Supplemental Figure 11. Relative proportions of each repeat category in relation to genome size.** Scatterplots showing the percentage of each repeat category from each species, plotted with respect to genome size. Statistical output from linear regressions shown for each repeat type.

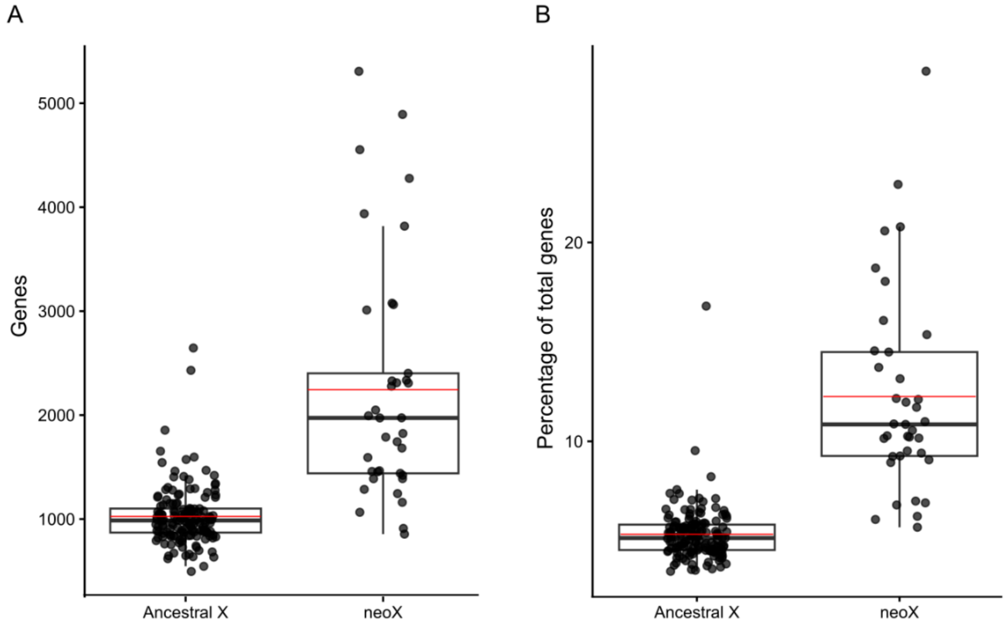


**Supplemental Figure 12. Comparison of gene content between ancestral X and neo-X chromosomes.** A) The number of protein-coding genes located on ancestral X chromosomes and inferred neo-X chromosomes (ancestral + fused chromosome(s)) of 190 species. B) The percentage of total genes located on the ancestral X or neo-X (ancestral + fused chromosome(s)). Percentages were calculated relative to the total number of annotated genes in each genome.
